# Atonosomes, compartments involved in membrane tension decrease

**DOI:** 10.64898/2026.07.03.736083

**Authors:** Elda Bauda, Aleksandr Aleksandrov, Maria Tettamanti, Julia Coronas Serna, Alice Gros, Nesli Sen, Margot Riggi, Paraskevi Linardou, Caroline Gabus, Geraldine Sylvano, Jean Daraspe, Sophie Martin, Omaya Dudin, Emmanuel Levy, Andreas Boland, Robbie Loewith

## Abstract

To ensure survival, cells need to buffer the effects of environmental stress on their plasma membrane, yet the structural mechanisms by which this is acutely achieved remain largely unknown. Here, we propose atonosomes as a unifying identity for a class of previously observed but enigmatic, tension-responsive, plasma membrane-derived compartments that arise across contexts of acute and chronic membrane tension loss. Leveraging unprecedented high resolution cryo-FIB-ET imaging in yeast, we show that atonosomes are complex, organelle-containing structures bounded by membranes and cell wall material, spanning hundreds of nanometers, and displaying a remarkable morphological diversity. Atonosomes form within seconds in response to reduced plasma membrane tension, and their emergence appears to require no dedicated molecular machinery, arising instead as a direct consequence of membrane biophysics. Upon formation, they recruit key membrane-associated proteins, including TORC2, Slm1, and septins. Under conditions of chronic disruption of PM homeostasis, atonosomes become constitutively present. Their stability and reversibility are further modulated by the cell wall, whose polymerization state influences atonosome dynamics. Structural conservation in fungi and ichthyosporea, demonstrates that atonosomes are a conserved stress-triggered response of cell-wall enclosed organisms. Together, these findings establish atonosomes as a novel compartment that mediates cellular responses to plasma membrane tension variation, coupling membrane remodeling and lipid homeostasis to preserve cellular integrity under stress.

## Introduction

Membrane compartmentalization is a critical strategy by which cells organize their functions. Rather than serving as homogeneous barriers, cellular membranes are structured into distinct domains that coordinate specific lipids and proteins, enabling the spatial coordination of signal transduction, interactions with the environment, and stress response ^1,2^. Over long timescales, unicellular organisms can tune plasma membrane (PM) lipid composition to optimize fitness for a given environment. In contrast, acute perturbation, including osmotic changes, are too rapid for lipid exchange. Instead, cells initially rely on phase transition and three-dimensional reorganization of pre-existing lipids to alleviate membrane stress^3,4^.

In yeast osmotic shocks directly affect PM domains such as eisosomes and key regulators of lipid homeostasis, including the Target of Rapamycin Complex 2 (TORC2). Eisosomes are stable structures primarily scaffolded by BAR domain proteins Pil1 and Lsp1 ^5–9^. TORC2 is a conserved protein kinase that regulates PM homeostasis through diverse processes influencing turgor pressure and lipid composition ^10,11^. Upon acute increase in PM tension caused by hypoosmotic shock, the essential paralogous proteins Slm1 and Slm2 are thought to exit eisosomes to activate TORC2 ^5^. Conversely, conditions that reduce PM tension, such as acute hyperosmotic stress or treatment with the small amphipatic molecule palmitoylcarnitine (PalmC), lead to TORC2 inhibition and the formation of membrane invaginations ^12^. These invaginations were originally described as phosphatidylinositol 4,5-bisphosphate (PI(4,5)P_2_)-Enriched Structures (PES) ^12^. Similarly, PI(4,5)P_2_ clustering, termed Large Eisosome-Derived Structures (LEDS) were reported under conditions of reduced PM tension ^13^, while related PM invaginations were also observed during hyperosmotic shock-induced desiccation ^14^. Comparable structures, referred to as large invaginations, "eisosome remnants" or "dimple-like structures" have further been described in *pil1Δ* ^9,15,16^ and in *inp51Δ inp52Δ* cells ^17–19^. Thus, PM structures have repeatedly appeared in the literature under diverse physiological and genetic contexts, often under different names. Despite their prevalence, they have not been systematically characterized, and high-resolution structural information is lacking.

Here, using cryo–focused ion beam milling and electron tomography (cryo-FIB-ET), we unify these observations and show that “PES”, “LEDS”, “eisosome remnants”, “dimple-like structures” represent a single, previously unrecognized compartment, distinct from eisosomes, that forms in response to acute loss of membrane tension or chronic disruption of PM homeostasis. We term these structures atonosomes (from Greek *a-* “no; without”, *tónos* “tension” and *sôma* “body”). With an unprecedented level of resolution, we show that atonosomes can span several hundred nanometers and form multilumen compartments bounded by two membranes and an intervening layer of cell wall (CW) material.

Acute-stress-induced atonosomes assemble within seconds following sudden decreases in PM tension (hyperosmotic shock or PalmC treatment), and their formation appears highly robust - a genome-wide screen failed to reveal any single gene deletion that would abolish their assembly. This remarkable plasticity suggests that membrane biophysics make a strong and direct contribution to their generation. Moreover, their appearance is tightly temporally coupled to TORC2 inactivation and recruitment, together with the accumulation of Slm1 and septins, indicating that atonosomes function as a signaling and assembly hub. In contrast, constitutive atonosomes are stable and reflect chronic disruption of membrane homeostasis. These structures are observed not only in *pil1Δ* and in *inp51Δinp52Δ* cells but also across a broad subset of deletion mutants affecting eisosomes, lipid synthesis and vesicular trafficking. Finally, the presence of atonosomes in the distantly related yeast *Schizosaccharomyces pombe* and the ichthyosporean *Chromosphaera perkinsii* demonstrate that they represent a conserved response to membrane tension loss in cell wall-enclosed organisms. Altogether, these findings unify previous cryptic observations to reveal a PM compartment arising under conditions of acute membrane tension loss or chronic disruption of tension homeostasis to likely function as a structural remodeling system and recruitment hub to maintain cellular integrity under stress.

## Results

### Cryo-FIB-ET of *S. cerevisiae* under acute PM tension decrease reveals atonosome formation and structure

Cryo-FIB-ET demonstrates that PM tension reduction, induced by either hyperosmotic shock or the membrane-intercalating agent PalmC, drives the formation of large compartments at the PM (**Fig.1, Movie S1**). We term these domains atonosomes derived from the Greek word tónosmeaning tension. Atonosomes are large, membrane structures with at least one lumen, spanning several hundred nanometers and displaying remarkable morphological diversity while sharing common features **(Fig. 1A-1C, S1A)**. The base of atonosomes originates at the PM, from which the compartment progressively invaginates into the cytoplasm to form a dome-shaped structure, with the tip located at its apex **(Fig. 1A)**. Atonosomes are separated from the cytoplasm by a layer of cell wall (CW) material, itself enclosed by an outer and an inner membrane (OM and IM); we refer to this enclosed space as the peripheral region **(Fig. 1A).** Peripheral regions vary in closure state: some appear fully connected to the cell envelope, isolating the atonosome lumen from the cytoplasm **(Figs. 1Bi, Ci–ii, S1B)**, whereas others are partially open **(Fig. 1Bii)**. In some instances, atonosomes remain connected to the PM only by a neck-like constriction at its base **(Fig. 1Ci)**. Both the OM and IM are closely associated with the cortical endoplasmic reticulum (ER) **(Figs. 1A, 1Bi, 1C, S1C)**. The lumen of atonosomes either contains amorphous material **(Fig. 1Bii, 1Ci)**, or includes identifiable cytoplasmic components such as the ER, ribosomes, Golgi elements and vesicles **(Fig. 1Ci, 1D)**. In addition, remarkable bundles of filamentous structures of unknown identity are occasionally observed within the lumen **(Fig. 1E)**. Statistical analysis of segmentation-derived dimension measurements demonstrates that atonosomes generated under hyperosmotic stress are significantly smaller and structurally less complex (x = 510 nm ± 220 nm, y = 560 nm ± 220 nm, average extrapolated volume 0.087 µm^3^; mean breadth: 1.5 ± 0.7 µm) than those formed under PalmC treatment (size: x =770 nm ± 300 nm, y= 830 nm ± 360 nm, average extrapolated volume 0.27 µm^3^; mean breath: 2.5 ± 1.1 µm) **(Fig. 1F)**. Mean breadth reflects the overall projected diameter of the compartment, which increases with number of cavities, reflecting complexity. Atonosomes are structurally distinct from much smaller eisosomes, which exhibit a characteristic canoe-like shape (Bharat et al., 2018, **Fig. 1G**).

**Figure 1.**
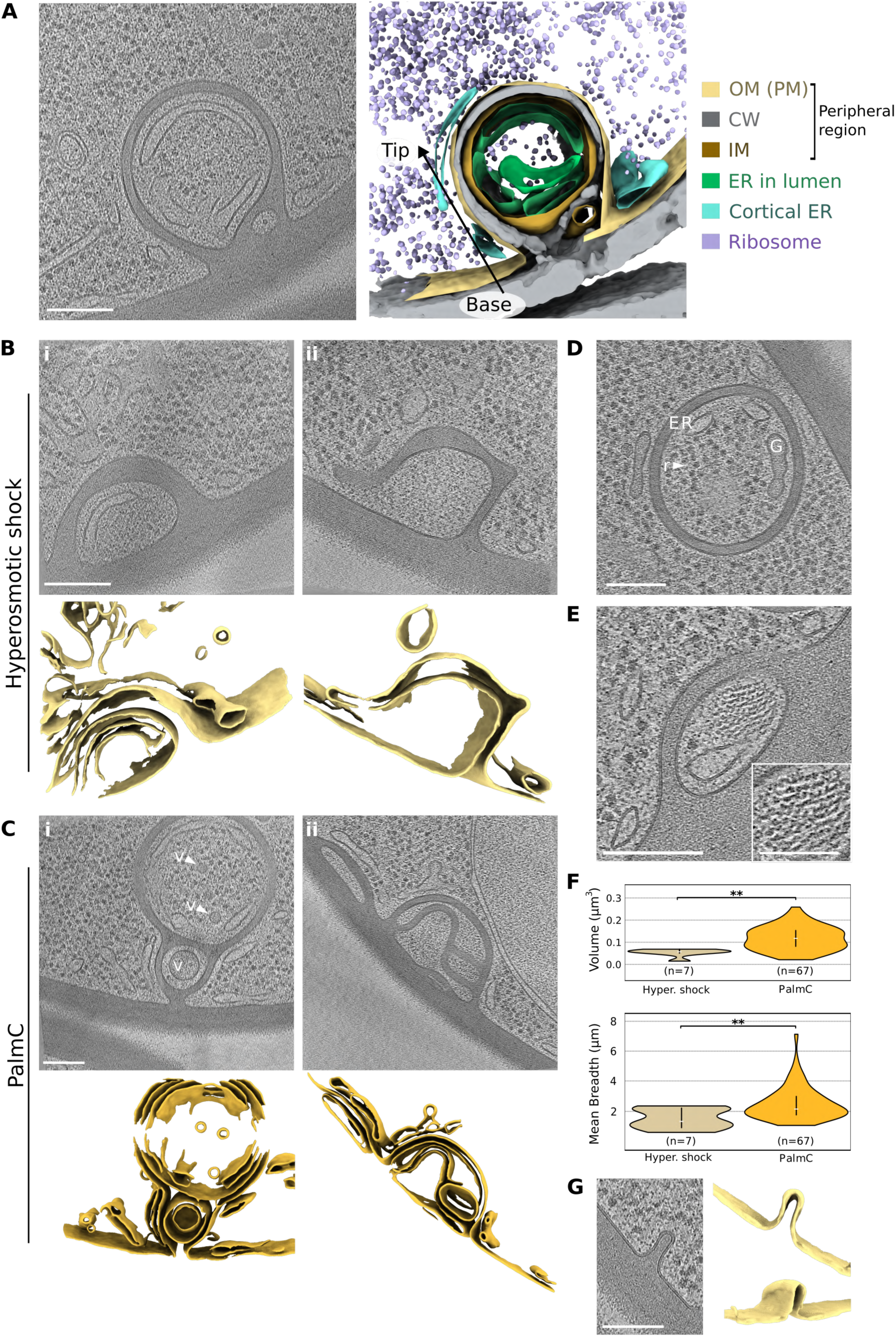
Cryo-electron tomography of *S. cerevisiae* under acute plasma membrane tension decrease reveals atonosome formation. **(A)** Cryo-tomographic slice and corresponding segmentation of a representative stress-induced atonosome in *S. cerevisiae.* The atonosome lumen, delimited by the inner membrane (IM), contains endoplasmic reticulum (ER) and ribosomes. Cortical ER is tightly associated with the outer membrane (OM) of the atonosome arising from the plasma membrane (PM). The IM and OM define a peripheral region containing cell wall material (CW). See Figure S1 for morphological diversity. **(B-C)** Cryo-tomographic slices and corresponding membrane segmentations of atonosomes formed under hyperosmotic shock **(B)** or Palme treatment **(C).** Compartments are either fully enclosed **(Bi, C)** or appear to be in the process of formation while remaining open to the cytoplasm **(Bii).** Some lumens contain cytoplasmic material including vesicles (*v*, arrows) **(Bi, Ci),** whereas others display amorphous content **(Bii, Cii)**. **(D)** Golgi apparatus (G), endoplasmic reticulum (ER), and ribosomes (r, arrow) are present within the atonosome lumen. **(E)** Bundles of filamentous structures of unknown identity within the atonosome lumen, with magnified view (scale bar, SO nm). **(F)** Segmentation-based quantification of atonosome volume and mean breadth, reflecting size in the lamellae and structural complexity, respectively."**" *p* value< 0.01; white horizontal bar, median; black vertical bar, IQR or middle 50%. **(G)** Eisosomes are observed at the plasma membrane and display a characteristic canoe-shaped morphology. Scale bars, 100 nm unless otherwise indicated.

Overall, atonosomes, previously reported in the literature under various names and experimental contexts, are in fact distinct structural entities with a defined lumen, organelle-like organization, and responsiveness to cellular stress.

### Atonosomes are dynamic, biophysically driven membrane compartments that act as a protein recruitment hub including Slm1, TORC2 and septins

To visualize atonosomes in vivo, we use fluorescent PI(4,5)P_2_- and phosphatidylserine- (PS) binding probes to label the PM ^21,22^. First, we performed cryo-CLEM to confirm that the observed fluorescent puncta formed under acute loss of PM tension indeed correspond to atonosomes observed in cryo-lamellae **(Fig. S2A)**. Having validated their identity, we next employed spinning-disk confocal live imaging to track their dynamics at 2–6 sec temporal resolution. Following hyperosmotic shock, cells rapidly shrink and form atonosomes within ∼5 s, with an average of 14 ± 4 atonosomes per cell (n=65) **(Fig. 2A, Movie S2a)**. In contrast, PalmC treatment induces the formation of 2 ± 1 atonosomes per cell approximately 1 min after application (n=65); these structures appear larger, consistent with cryo-ET observations **(Fig. 2B, Movie S2b**). Together, these observations reveal that atonosomes are dynamic membrane compartments characterized by the accumulation of membrane (and cell wall material) rather than by enrichment of a single, defining lipid species. Consequently, their previous designation as “PI(4,5)P₂-Enriched Structures” (PES) ^12^ does not fully capture their composition or behavior. Moreover, recent work has further shown that sterols are also transiently enriched at atonosome sites in *S. cerevisiae* ^23^.

**Figure 2.**
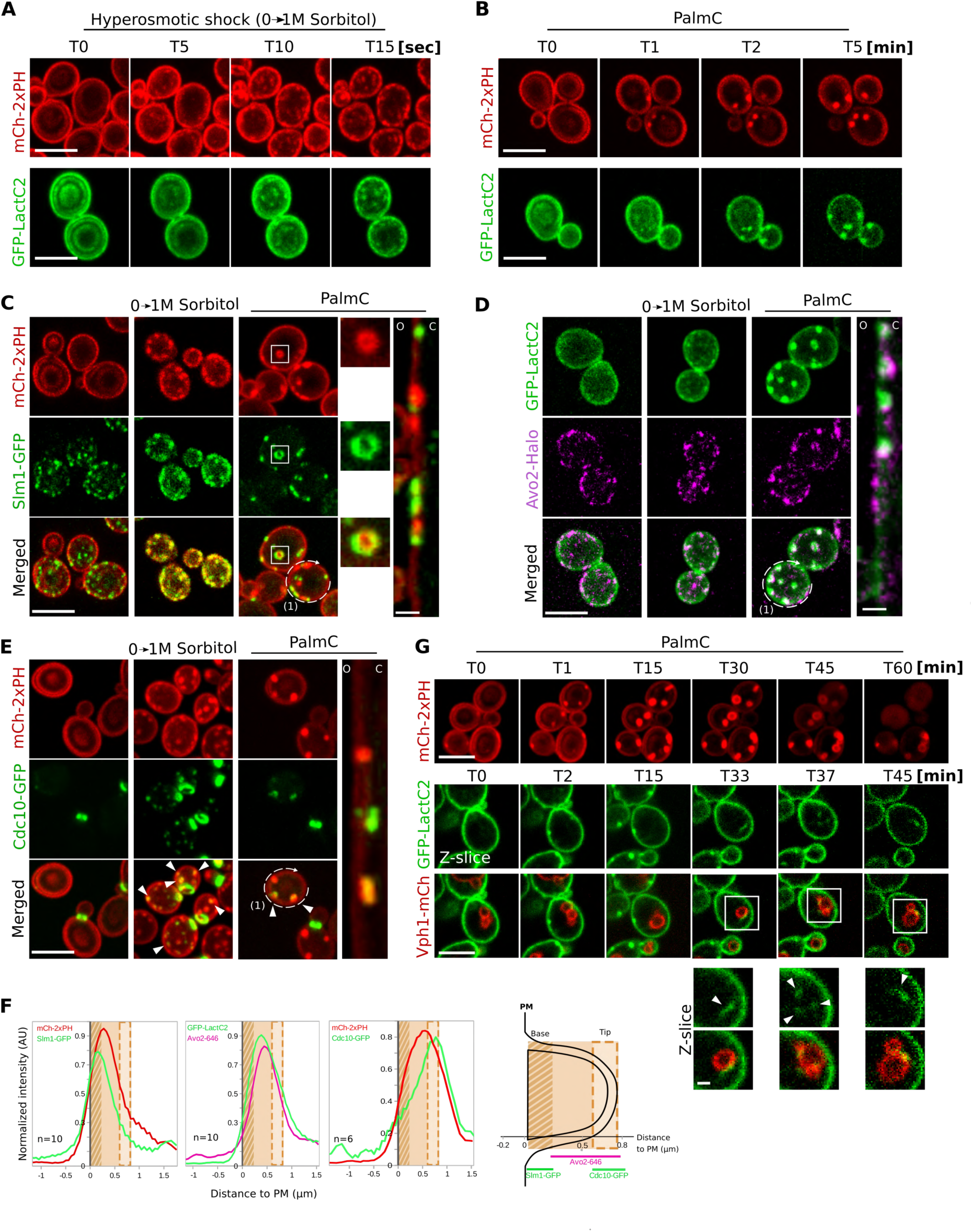
atonosomes are dynamic, biophysically driven membrane compartments that act as a protein recruitment hub including Slm1, TORC2 and septins. **(A-B)** Live spinning-disk confocal microscopy showing rapid atonosome formation following hyperosmotic shock **(A)** or Palme treatment **(B).** The plasma membrane is visualized in *WT* cells expressing an endogenous Pl(4,5)P_2_ reporter (meh-2xPH) or a PS reporter (GFP-LactC2) on a plasmid. **(C)** Spinning-disk confocal microscopy of cells co-expressing endogenously Slm1-GFP and the Pl(4,5)P_2_ reporter (meh-2xPH). The outer right panel shows a representative linearized fluorescence profile generated along (1) the cell periphery (scale bar, 1 µm). O, outside the cell. C, cytoplasm. **(D)** Spinning-disk confocal microscopy of cells expressing a PS reporter (GFP-Lacte2) from a plasmid and endogenous Avo2-Halo (TORC2 subunit) labeled with fluorescent halo ligand 646. The outer right panel shows a representative linearized fluorescence profile generated along (1) the cell periphery (scale bar, 1 µm). O, outside the cell. C, cytoplasm. **(E)** Spinning-disk confocal microscopy of cells expressing endogenous septin edc10-GFP and the Pl(4,5)P_2_ reporter. The outer right panel shows a representative linearized fluorescence profile generated along (1) the cell periphery (scale bar, 1 µm). O, outside the cell. C, cytoplasm. **(F)** Averaged fluorescence intensity profiles and schematics of Slm1-GFP, Avo2-Halo, and edc10-GFP positioning, measured perpendicular to atonosomes identified with Pl(4,5)P_2_ or PS reporters. The relative positions of fluorescence peaks indicate the localization of each protein with respect to the PM, the atonosome base and tip. **(G)** Live spinning-disk confocal microscopy showing the fate of Palme-induced atonosomes over long time periods. Puncta of the PS reporter (GFP-Lacte2) (white arrows) are seen at the vacuole surface (Vph1-meh) around 30 minutes after Palme treatment. Insets scale bar, 1 µm. For all panels except live imaging, hyperosmotic shock was induced with 1 M sorbitol in culture medium for 1 min. Palme treatments were performed at 8 µM for 5-10 min. Unless otherwise indicated, all images are maximum-intensity projections along the z-axis and scale bars, 5 µm

While the characteristic canoe-shape of eisosome can be clearly distinguished from atonosomes morphology by cryo-ET **(Fig. 1G)**, atonosome triggered by hyperosmotic shock was previously described as “Large Eisosome Derived Structures” (LEDS), arguing that eisosome could transform into such structures upon PM tension loss ^13^. These observations prompted us to further investigate the relationship between the two structures. Imaging of Pil1–GFP in a *WT* background shows no consistent colocalization between Pil1 and atonosomes **(Fig. S2B).** Furthermore, atonosomes were frequently observed in buds lacking detectable Pil1–GFP signal **(Fig. S2B**), consistent with previous reports showing that eisosomes are generally absent from buds ^24^. In contrast, the eisosomal protein Slm1 dynamically relocates to the compartments, often forming enriched horseshoe-like patterns around their base **(Fig. 2C, 2F)**. Together, these findings indicate that although Slm1 relocates to atonosomes, their formation is independent of eisosomes.

In addition to inducing atonosome formation, PM tension decrease caused by hyperosmotic shock or PalmC treatment also inhibits TORC2 (**Fig. S2C,**^12^) leading us to examine TORC2 localization. We confirm that TORC2 inactivation occurs concurrently with atonosome formation and is accompanied by relocation of TORC2 components (Avo2 and Avo3) to these compartments, with enrichment away from the base **(Fig. 2D, 2F, S2)**. Strikingly, cryo-electron tomograms revealed linear protein assemblies on the OM of atonosomes, reminiscent of septin filaments at the bud neck **(Fig. S2E)**. The septin Cdc10 was indeed partially recruited, with septin signal redistributing from early division sites to newly forming atonosome tips **(Fig. 2E-F, Movie S3a,b)**. Overall, atonosomes act as recruitment hubs for membrane proteins such as Slm1, TORC2, and Cdc10, which partition into distinct regions within the compartment.

The rapid formation of atonosomes prompted us to test which PM organization factors would promote their assembly. Using PalmC-induced atonosomes for precise temporal control, we found that deletion of six ER–PM tethering proteins (t*cb1/2/3Δ scs2/22Δ ist2Δ*), which abolishes cortical ER, did not impair atonosome formation despite their apparent ER association in tomograms **(Fig. S2F)**. Similarly, although atonosomes sometimes display neck-like constrictions, deletion of key ESCRT components (*vps4Δ, vps25Δ, did4Δ*) had no effect **(Fig. S2G)**. Notably, a genome-wide screen of >5,000 deletion strains failed to identify mutants with severe defects in atonosome formation (data not shown), suggesting that their rapid emergence does not require dedicated machinery but instead arises from biophysical changes in PM properties. This observation is compatible with previous evidence showing that atonosome formation was not energy driven and did not require *de novo* synthesis of lipids ^12^. Overall, it appears that atonosome genesis is independent of eisosomes and active cellular processes, pointing to a mechanism driven by biophysical forces alone.

An important question raised by the formation of these large membrane structures concerns their fate: are atonosomes recycled, or do they persist under sustained stress conditions? Within a few minutes following hyperosmotic shock, atonosomes become progressively less defined, and all compartments disappear after 25 minutes **(Fig. S3A)**, consistent with previous kinetics measurements ^13^. Following PalmC treatment, some atonosomes disappear after 30 minutes **(Figs. 2G, S3Bi–ii)**, whereas others remain associated with the PM for up to 2 hours, the longest time point examined **(Fig. S3Biii)**. During this period, the PI(4,5)P₂ probe sometimes dissociates from the PM, likely reflecting PI(4,5)P₂ dephosphorylation or hydrolysis **(Fig. S3Bi)**. In contrast, when the PS probe is used, PS-positive puncta appear at the vacuole membrane following the disappearance of atonosomes from the PM **(Fig. 2G)**. In some instances, we observe formation and extension of atonosomes into the cytoplasm before PS-positive puncta detach and migrate toward the vacuolar surface **(Fig. S3C)**. These observations suggest that intact atonosomes may be trafficked to the vacuole for degradation. Building on our previous observations pointing to a critical role for retrograde transport of ergosterol in the context of acute membrane tension loss ^23^, dissection of the molecular mechanisms governing atonosome fate are ongoing.

Altogether, atonosomes emerge rapidly upon PM tension loss through a mechanism that does not require dedicated cellular machinery but mainly relies on biophysics. They function as dynamic recruitment hubs for signaling and structural proteins, including Slm1, TORC2, and septins. Atonosomes exhibit distinct fates, either disappearing or persisting, and may be targeted to the vacuole under sustained membrane excess.

### High-resolution imaging of eisosome mutants shows constitutive atonosome formation

TEM of resin sections and scanning EM of *pil1Δ* yeast cells under steady-state conditions uncovered striking aberrant membrane structures, termed "eisosome remnants", "dimple-like structures" or “large cell wall invaginations” ^9,15,16^. Atonosomes strongly resemble such structures. Using TEM of resin sections, we confirm that *pil1Δlsp1Δ S. cerevisiae* cells indeed exhibit constitutive atonosome-like structures **(Fig. S4A)**. We extended our study using high resolution cryo-FIB-ET of cells expressing a Pil1 variant that is defective in sterol binding and fails to localize to defined membrane domains (*pil1-sterol lsp1Δ*) ^7^. Cryo-FIB-ET further confirms that the architecture of these compartments closely resembles that of stress-induced atonosomes, with an astonishing diversity of morphology **(Figs. 3, S4B, Movie S4)**. These compartments, termed constitutive atonosomes due to their presence in steady-state growth conditions, can present multiple lumens, filled with cytoplasmic material **(Fig. 3A-Ci)** or with amorphous material **(Fig. 3Cii)**. The lumen is separated from the surrounding cytoplasm by the peripheral region (IM and OM enclosing CW material). Interestingly, bundles of filaments with unknown identity are also detected in the lumen **(Fig. 3Di-ii)**. Segmentation-based measurements show that constitutive atonosomes have a similar size and complexity as observed for PalmC-induced counterparts (x = 902 nm ± 372 nm, y = 946 nm ± 392 nm, average extrapolated volume 0.4 µm^3^; mean breadth: 3.2 ± 1.8 µm) **(Fig. 3E).**

**Figure 3.**
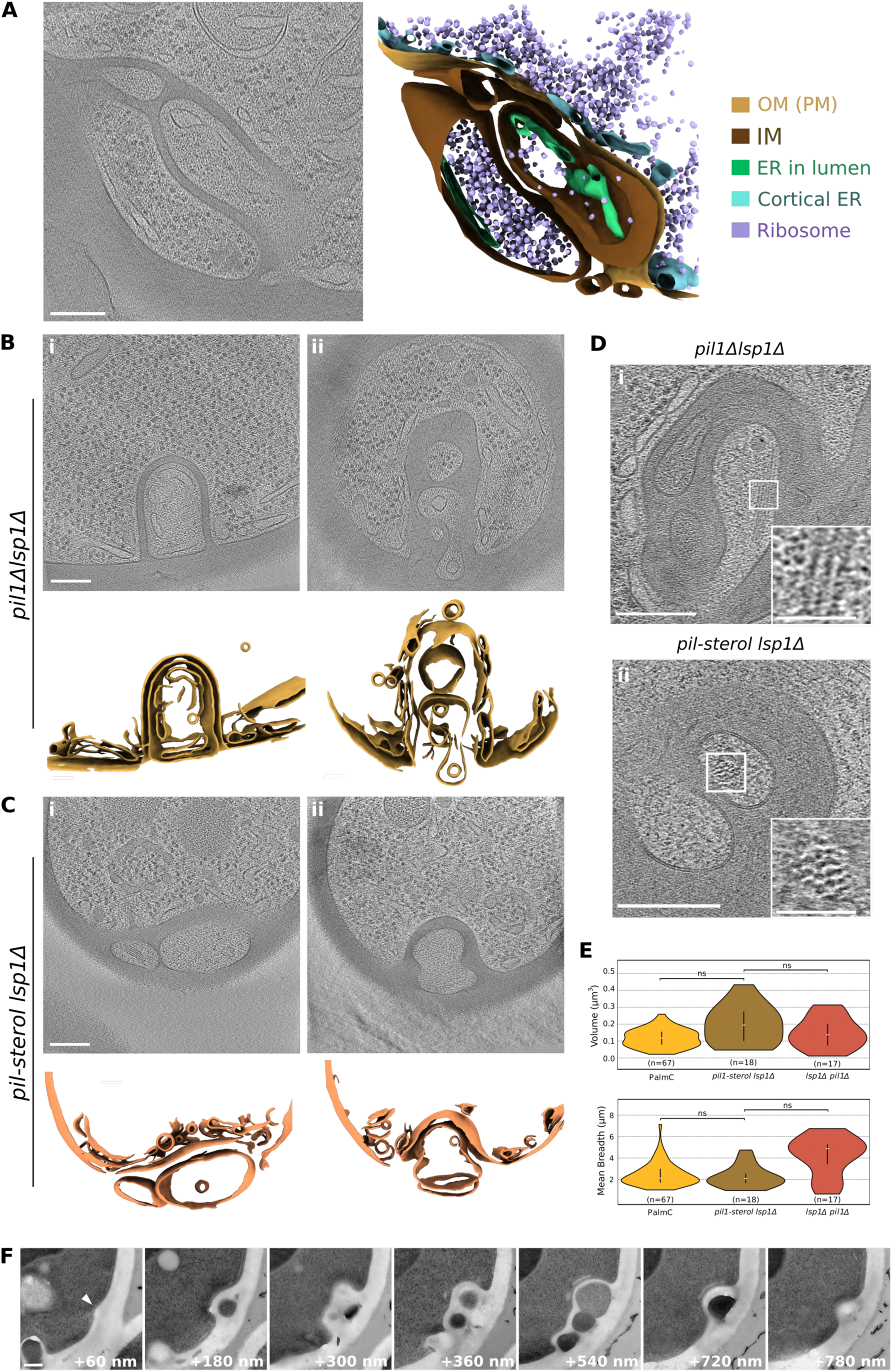
High-resolution imaging of eisosome mutants reveals constitutive atonosome formation. **(A)** Cryo-tomographic slice and corresponding segmentation of a constitutive atonosome in *pil1Δ lsp1Δ S.cerevisiae* cells. The atonosome lumen, delimited by the inner membrane (IM). contains endoplasmic reticulum (ER) and ribosomes. Cortical ER is tightly associated with the outer membrane (OM) of the atonosome arising from the plasma membrane (PM). The IM and OM define a peripheral region containing cell wall material. See Figure S2 for morphological diversity. **(B-C)** Cryo-tomographic slices and corresponding membrane segmentations of atonosomes in *pil1Δ lsp1Δ* **(B)** or *pi/1-stero/ /spli1* **(C)** cells. The lumen of some compartments contains organelles and ribosomes **(Bi-ii, Ci),** whereas others display amorphous content **(Cii).** **(D)** Bundles of filamentous structures of unknown identity within the atonosome lumen, with magnified views (scale bar, 50 nm) from *pil1Δ lsp1Δ* **(i)** and *pill-sterol lsp1Δ* cells **(ii)**. **(E)** Segmentation-based quantification of constitutive versus PalmC-induced atonosome volume and mean breadth, reflecting size and structural complexity, respectively. "ns", not signifiant. **(F)** Serial sectioning of resin-embedded *pil1Δ/spliΔ.* cells showing sequential slices through an atonosome volume. Numbers indicate relative thickness with respect to the first section on which atonosome appears. Scale bars, 100 nm unless otherwise indicated

Cryo-FIB-ET, in which 150 nm-thick sections are produced and imaged, typically do not contain an entire atonosome. To overcome this challenge, we implemented serial sectioning on resin sample. Consistently with our cryo-FIB-ET observations, serial sections of an entire atonosome volume revealed that they can exceed 800 nm in length, with some compartments spanning over 15 consecutive 60 nm-thick sections. The presence of CW material at both extremities, and along its base and tip, shows that constitutive atonosomes can become fully enclosed, with their lumen being isolated from the cytoplasm and the external environment **(Fig. 3F, Movie S5)**.

Constitutive atonosomes can be visualized by fluorescence microscopy using a PI(4,5)P₂ reporter in *pil1Δ lsp1Δ* and *pil1-sterol lsp1Δ* cells **(Fig. S4C)**. To assess the role of tetraspanners in constitutive atonosome formation, we used a mutant lacking the six major tetraspanin superfamily proteins in addition to *pil1Δ* (*6tspΔ: sur7Δ, fmp45Δ, ynl194cΔ, pun1Δ, nce102Δ, fhn1Δ*) ^15^. In this background as well, atonosomes remain constitutively present at the PM **(Fig. S4C)**. As observed upon acute loss of membrane tension, Slm1 and septins associate with constitutive atonosomes, with Slm1 forming protein-rich structures and Cdc10 localizing as stable foci **(Fig. S4D,Ei)**; consistent with this association, septin-like filaments are also detected on the OM by cryo-FIB-ET **(Fig. S4Eii)**. Interestingly, deletion of *SLM1/2* in addition to all major eisososome components (*pil1Δ slm1Δ slm2Δ 6tspΔ)* completely abolishes the constitutive atonosome phenotype: the PM appears smooth and uniform **(Fig. S4G “T0”)**. The implications of this latter result are explored below.

We next asked whether constitutive atonosomes remain responsive to acute loss of PM tension. Under hyperosmotic shock, they are rapidly perturbed: the compartments become highly dynamic, showing apparent movement and shrinkage, but remain detectable several minutes after the shock **(Fig. S4F, Movie S6a)**. Upon PalmC treatment, constitutive atonosomes initially decrease in size, but within one minute drastically expand, in some cases extending across the entire cytoplasm **(Fig. S4F, Movie S6b)**. Interestingly, the *pil1Δ slm1Δ slm2Δ 6tspΔ* mutant, in which constitutive atonosomes are absent, show stress-induced compartment formation with kinetics comparable to WT **(Fig. S4G, Movie S7a,b)**, further supporting that atonosome formation is independent from eisosome.

In conclusion*, pil1Δ lsp1Δ, pil1-sterol lsp1Δ,* and *pil1Δ lsp1Δ 6tspΔ* mutants form constitutive atonosomes at the PM under steady-state conditions, whereas deletion of *SLM1/2* abolishes this phenotype. Constitutive atonosomes closely resemble stress-induced ones in their architecture, lumenal content, size, and protein composition.

### Constitutive atonosomes result from impaired lipid homeostasis

Slm1/2 are TORC2 activators normally sequestered at eisosomes ^5^. We therefore asked whether TORC2 signaling is altered in eisosome mutants. Upon acute loss of PM tension, *pil1Δ lsp1Δ, pil1-sterol lsp1Δ*, and, more strongly, *pil1Δ 6tspΔ* cells fail to properly inactivate TORC2, suggesting that loss of Slm1 sequestration leads to TORC2 dysregulation **(Fig. 4A)**. Consistent with this, *pil1Δ slm1Δ slm2Δ 6tspΔ* cells partially restore normal TORC2 inactivation **(Fig. 4A)**. We therefore hypothesized that TORC2 hyperactivation–like state, caused by absence of Slm1 sequestration at eisosomes, drives atonosome appearance in eisosome mutants. Consistent with this model, expression of a constitutively active Ypk1 variant (Ypk1-D242A), which mimics TORC2 hyperactivation ^25^, is sufficient to generate extensive compartments **(Fig. 4B)**, supporting a causal link between TORC2 hyperactivity and constitutive atonosomes.

**Figure 4.**
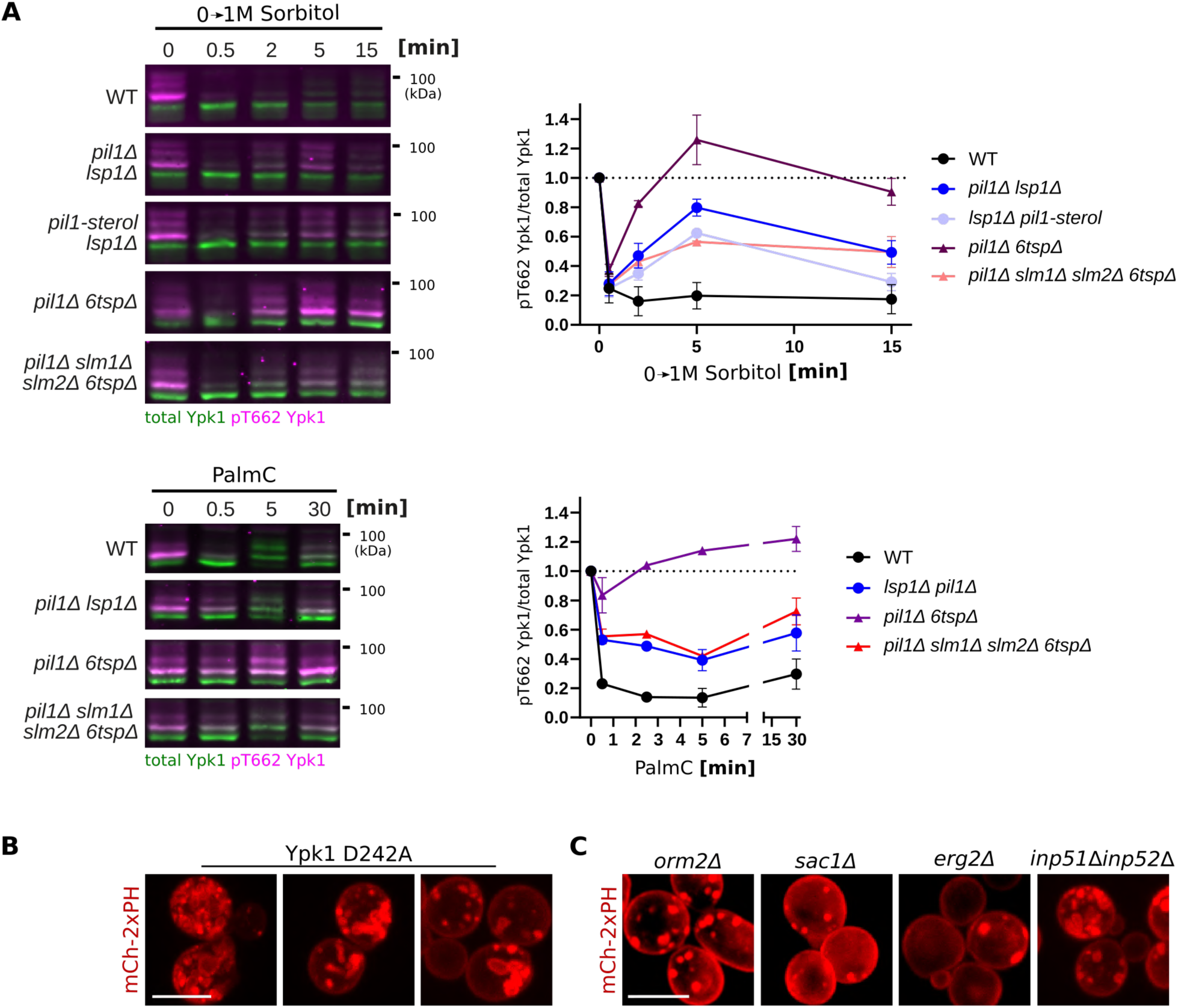
Constitutive atonosomes result from impaired lipid homeostasis. **(A)** Western blot analysis of TORC2 activity in wild-type (WT) cells and the indicated mutants following treatment with 1 M sorbitol or 5 µM Palme. TORC2 activity was assessed by relative phosphorylation levels of Ypk1. Quantification represents mean± s.d. from N = 3 independent experiments. **(B)** Spinning-disk confocal microscopy of cells expressing a Pl(4,5)P_2_ reporter (mCh-2xPH) endogenously and a constitutively active Ypk1 variant (Ypk1D242A) from a plasmid. Scale bar, 5 µm. **(C)** Spinning-disk confocal microscopy of deletion mutants involved in lipid homeostasis expressing a Pl(4,5)P_2_ reporter (mCh-2xPH) from a plasmid. Scale bar, 5 µm. All fluorescence images are maximum-intensity projections along the z-axis

To gain a broader, unbiased understanding of factors influencing PM homeostasis, we asked whether other genetic perturbations might also trigger constitutive atonosome formation. To this end, we screened a library of ∼5,000 deletion and essential knockdown mutants for membrane defects using PI(4,5)P_2_ and PS fluorescent probes imaged by confocal microscopy ^26,27^. By comparing the PM morphology, we identified 30 mutants that chronically present lipid clustering at the PM, reminiscent of atonosomes **(Table 1).** The screen also identified mutants with diffuse cytoplasmic PI(4,5)P_2_ signal. These deletions are not discussed here but are listed in **Table S1**. Among the hits showing atonosome-like elements, seven were deletions of genes related to lipid metabolism. Most notable among them were *orm2Δ, sac1Δ* and *erg2Δ*, which we further confirmed in a different *S. cerevisiae* background **(Fig. 4C)**. Orm2 and Sac1 are direct inhibitors of the SPT complex (serine palmitoyltransferase), which catalyzes the first step of de novo synthesis of sphingolipids ^28,29^. Besides interacting with the SPT complex, Sac1 also dephosphorylates phosphatidylinositol 4-phosphate (PI4P), thereby tightly restraining the PI levels ^30,31^. Interestingly, the deletion of other conserved PI phosphatases Inp51 and Inp52, involved in dephosphorylation of PI(4,5)P_2_ into PI4P ^19^, also result in constitutive atonosome formation **(Fig. 4C)**, consistent with large membrane and cell wall depositions previously observed by EM of resin section ^16,18,19^. Erg2 converts fecosterol into episterol in the ergosterol biosynthesis pathway ^32^. The deletion of the non-essential enzymes within the pathway leads to sterol intermediates accumulation and disordered PM ^33,34^.

**Table 1.**
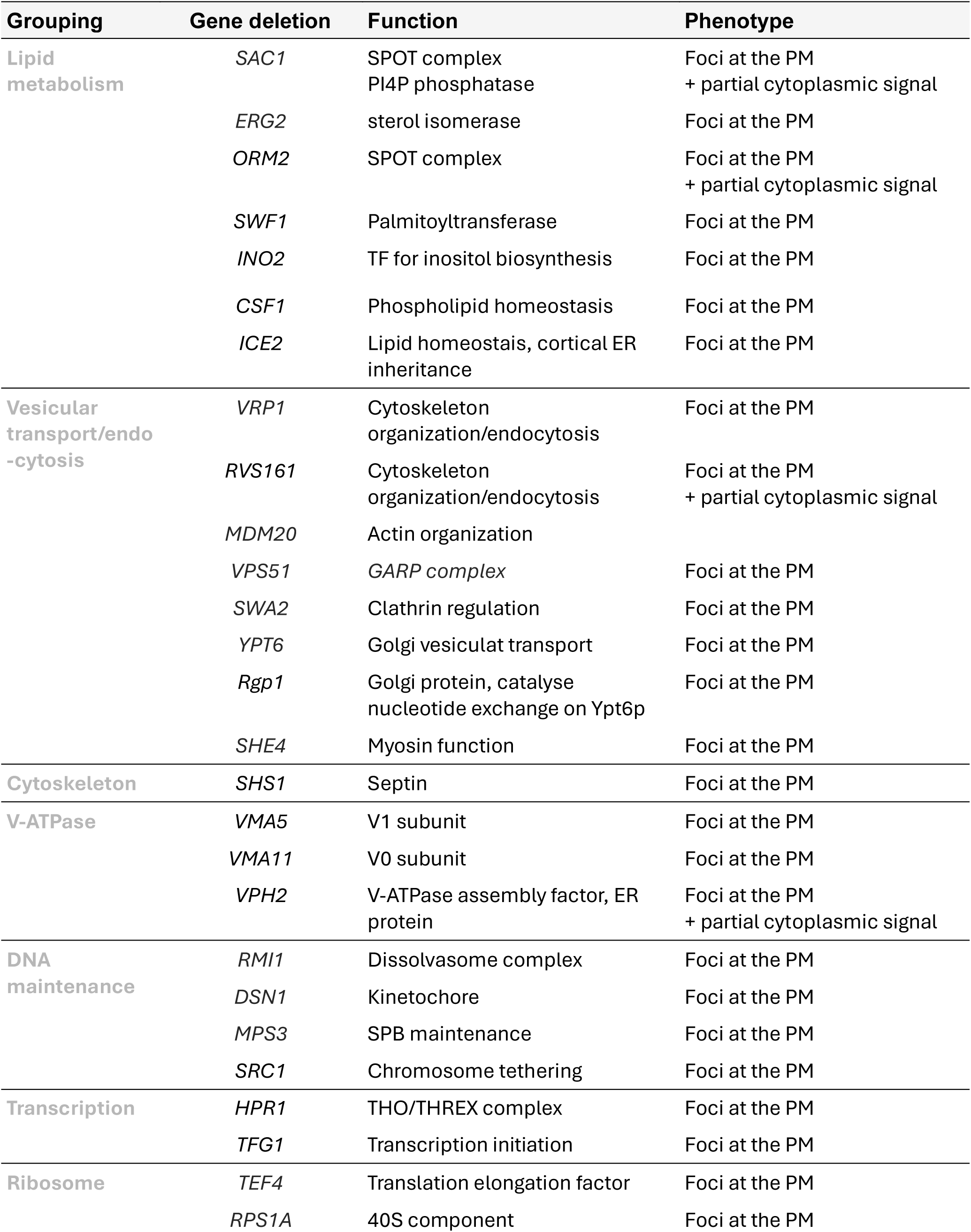
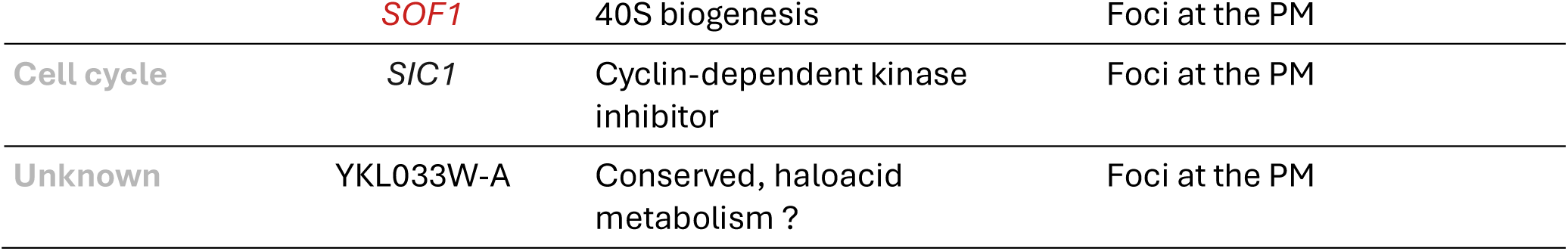
yMS2085 MATα strains carrying either a deletion of a single non-essential gene or downregulation of an essential gene (highlighted in red). All strains endogenously express fluorescent markers for PI(4,5)P₂ and phosphatidylserine (PS). Following confocal microscopy screening for plasma membrane (PM) defects, 30 strains exhibited fluorescent lipid clustering (foci) at the PM, reminiscent of atonosome phenotype. In some cells, lipid clustering at the PM coexisted with cytoplasmic fluorescence.

Together, these findings identify impaired lipid homeostasis, arising either from TORC2 hyperactivity in eisosome mutants or from disrupted lipid synthesis in *orm2Δ*, *sac1Δ*, *erg2Δ*, and *inp51*Δ inp52Δ, as the driver of constitutive atonosome formation.

### The cell wall contributes to atonosome architecture and dynamics

Cryo-ET reveals the ultrastructure of atonosomes, the PM and the CW at nanometric resolution. The CW appears as a diffuse layer with dark gray density, continuous from the hair-like outer layer to the PM surface, itself delineated by two dark lines corresponding to the head groups of the phospholipid bilayer **(Fig. 5A)**. A density, identical to the one of the CW, is present in the peripheral region of atonosomes **(Fig. 5A)**. Interestingly, granular darker densities are observed within the constitutive atonosome peripheral regions **(Fig. 5A)**. The apparent presence of CW material in such a dynamic structure remains puzzling, as CW polymers are generally considered to be rigid and non-deformable ^35–37^. Therefore, to further confirm that the peripheral region of atonosomes contains CW material, we assessed calcofluor white (CFW) stained cells in which chitin and β(1,3)-glucan fibers are fluorescently labelled ^38,39^. Interestingly, stress-induced atonosomes do not display any calcofluor white staining, while constitutive atonosomes are robustly labelled **(Fig. 5B)**. We examined whether these staining differences reflect distinct CW polymerization states by observing resorption dynamics of stress-induced versus constitutive atonosomes. Notably, stress-induced atonosomes rapidly disappear upon subsequent hypoosmotic shock, whereas constitutive atonosomes remain stable and largely immobile **(Fig. 5C, Movie S8a,b,c,d)**.

**Figure 5.**
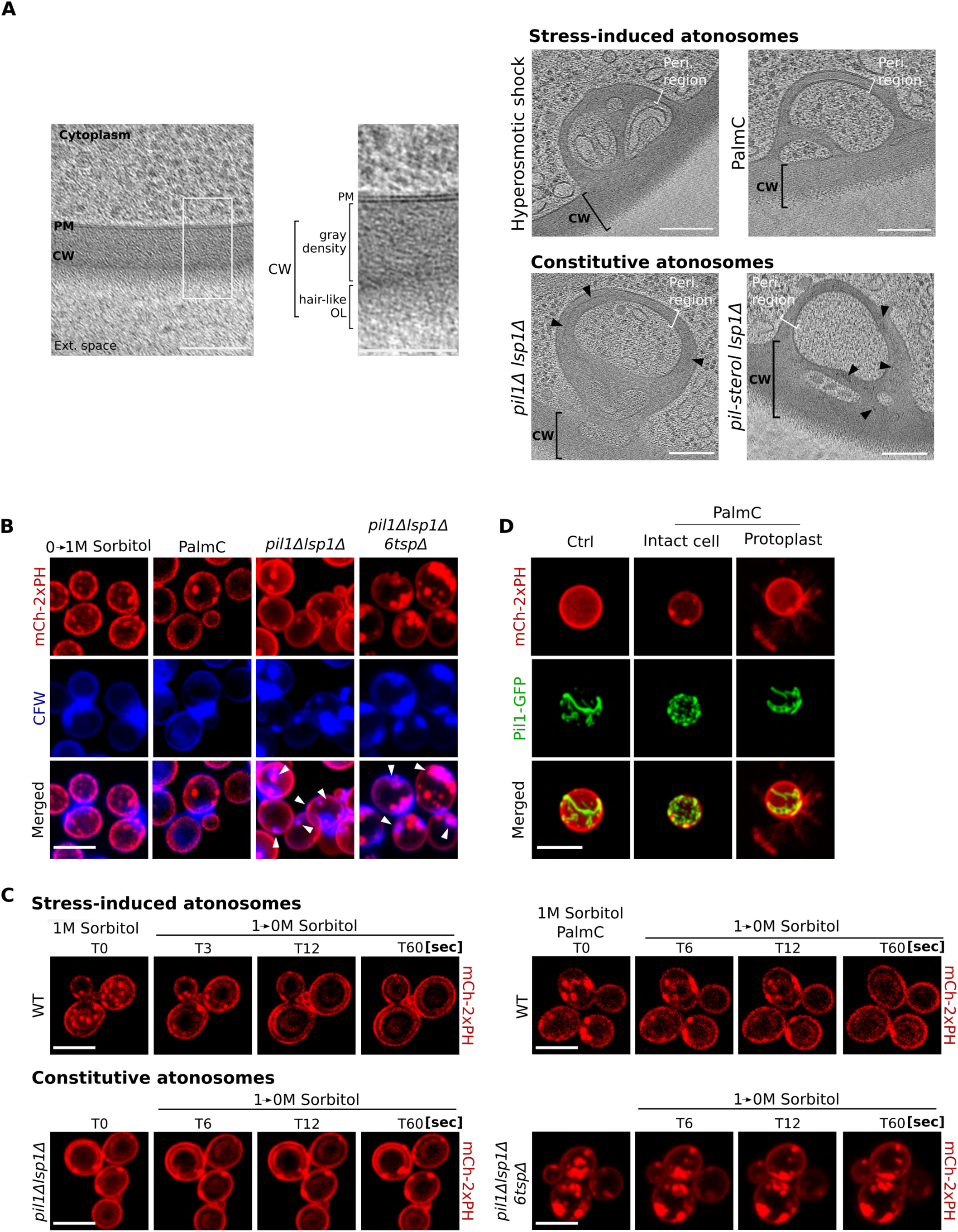
The cell wall contributes to atonosome architecture and dynamics. **(A)** Cryo-tomographic slice showing the representative architecture of the plasma membrane (PM), the cell wall (CW) and of the peripheral region of stress-induced and in constitutive atonosomes. The CW is characterized by a gray density and a hairy-like outer layer (OL). Black arrows in peripheral regions of constitutive atonosomes show granular aspect of CW material. Scale bar, 100 nm. **(B)** Spinning-disk confocal microscopy of WT cells under hyperosmotic shock (1M sorbitol for 1 min) or Palme treatment (8 µM for 5 min) and indicated eisosome deletion mutants expressing Pl(4,5)P_2_ reporter (mCh-2xPH) endogenously. Cells were stained with calcofluor white (CFW). White arrows show colocalization between CFW and mCh-2xPH signals. **(C)** Live spinning-disk confocal microscopy of atonosomes under osmotic stress. Atonosome formation was induced by acute hyperosmotic shock (1 M sorbitol, 1 min) or Palme (8 µM in culture medium with 1M sorbitol for 5 min) in cells pre used to high osmotic medium. For hypoosmotic shock, cells were pre-adapted to 1 M sorbitol and then shifted to sorbitol-free medium. Constitutive atonosomes in eisosome-deficient mutants were subjected to identical conditions. **(D)** Spinning-disk confocal microscopy of protoplasts of WT cells co-expressing endogenously a Pl(4,5)P_2_ reporter (mCh-2xPH) and Pil1-GFP, subjected to Palme (8 µM in culture medium with 1M sorbitol For 5 min). A non protoplasted intact cell is shown for comparison. All fluorescence images are maximum-intensity projections along the z-axis. Scale bars, 5 µm unless otherwise indicated.

We then asked whether CW material plays a structural role in atonosome genesis or structure. To this end, we studied PalmC response in protoplasts – yeast cells devoid of a CW – using live confocal microscopy. As shown before, protoplasts present collapsed eisosomes that form long tubes at the cell surface **(Fig. 5D**, ^5^). Strikingly, in the absence of a CW, PalmC promotes the formation of long, highly mobile membrane protrusions extending outward **(Fig. 5D, Movie S9)**. Overall, variations in the aspect in tomograms, in calcofluor white staining and in the resorption, dynamics reflect different polymerization states in the CW, which would be more gel-like and flexible in acutely forming atonosomes, and more rigid and fibrous in constitutive compartments. The protrusions emerging from the PM in protoplasts upon PalmC treatment clearly demonstrate that the CW is essential for formation of inward atonosome architecture.

### Atonosomes are a conserved response to acute PM tension loss across cell wall- enclosed organisms

We wondered whether atonosomes would represent a conserved cellular response to membrane stress. To address this question, we investigated PM architecture in *Schizosaccharomyces pombe*, a distantly related ascomycete yeast that diverged from budding yeast more than 300 million years ago ^40^. Interestingly, using lipid reporter probes specific for PS, sterol^41^ and PI4P ^42^, we found that atonosomes form in response to PM tension decrease **(Fig. S5A)**. Their formation dynamics are identical to the ones in *S. cerevisiae*: atonosomes are seen ∼5 sec after hyperosmotic shock and ∼60 sec upon PalmC addition **(Fig. S5B-C, Movie S10a,b)**. In our hands, the PI(4,5)P_2_ probe scarcely stained the PM and did not highlight any atonosomes in similar conditions (data not shown).

Cryo-FIB-ET strikingly reveals that, both under hyperosmotic shock and under PalmC treatment, *S. pombe* atonosomes adopt a structure indistinguishable from *S. cerevisiae* atonosomes: they emerge from the PM, with their lumen separated from the cytoplasm by the peripheral region **(Figs. 6, S5D, MovieS11)**. Cytoplasmic content with recognizable organelles is present within the lumen (ER, ribosomes) **(Fig. 6A-C)**, and unknown bundles of filamentous structures are also detected **(Fig. 6D)**. Strikingly, we could also find septin-like filaments on the OM, further supporting conserved protein recruitment properties in both species **(Fig. S5F)**. Analysis of segmentation-derived volumes indicates that *S. pombe* atonosomes are similarly sized and complex under both stresses but significantly larger than those in *S. cerevisiae* **(Figs. 6E, S5E).** Atonosome’s architecture is distinct from eisosomes which also exhibit a canoe shape in *S. pombe* **(Fig. 6F)**.

**Figure 6.**
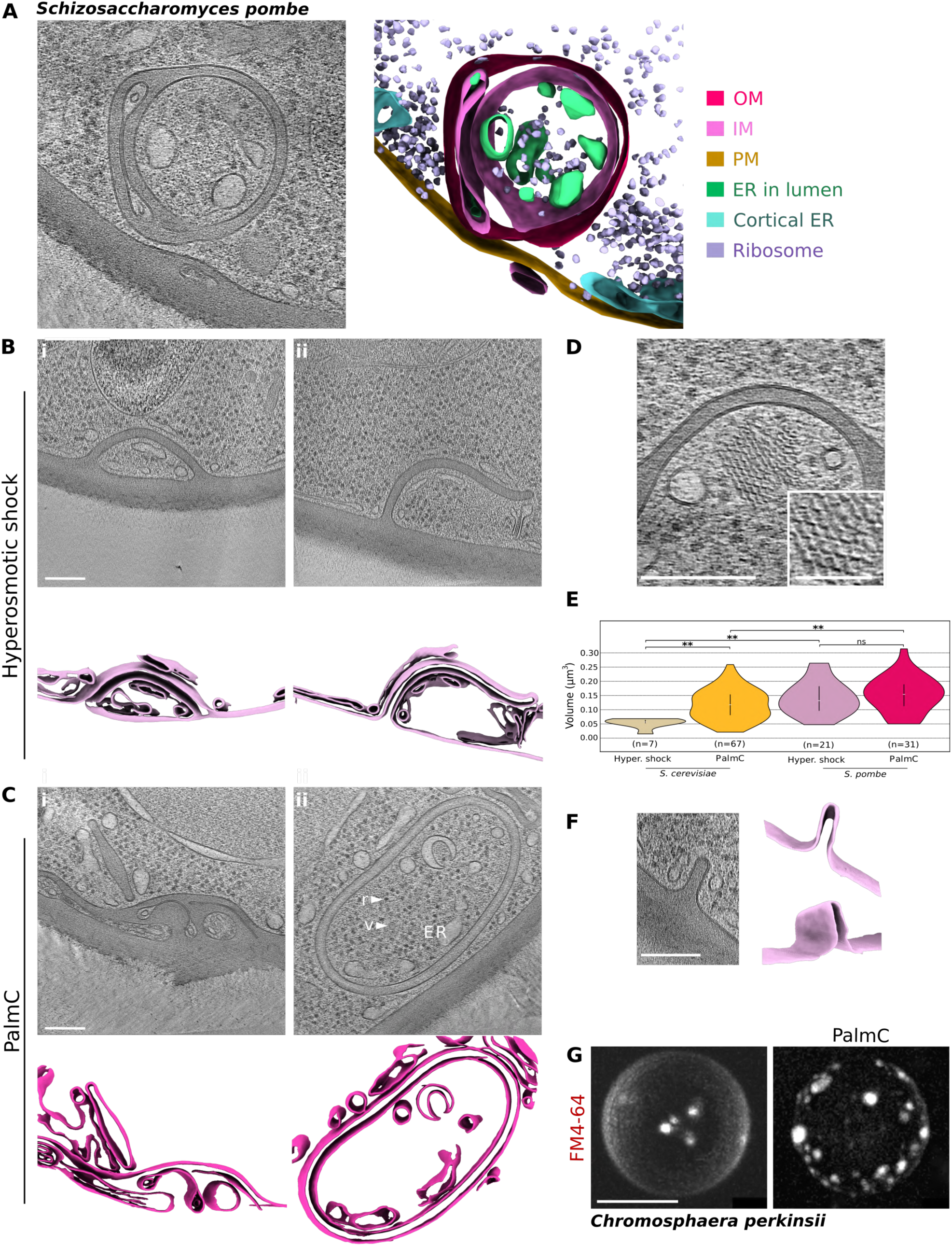
High-resolution imaging of atonosomes in *S. pombe* and *C. perkinsii*. **(A)** Cryo-tomographic slice and corresponding segmentation of a representative stress-induced atonosome in *S. pombe.* The atonosome lumen, delimited by the inner membrane (IM), contains endoplasmic reticulum (ER) and ribosomes. The IM and the outer membrane (OM) define a peripheral region containing cell wall material. See Figure S5 for morphological diversity. **(B-C)** Cryo-tomographic slices and corresponding membrane segmentations of stress-induced atonosome formed under hyperosmotic shock **(B)** or Palme treatment **(C).** Compartments are either fully enclosed **(Bi, C)** or appear to be in the process of formation while remaining open to the cytoplasm (Bii). The lumens contain a variety of organelles such as ER, vesicles (v, arrow), and ribosomes (r, arrow)(Cii). **(D)** Bundles of filamentous structures of unknown identity within the atonosome lumen, with magnified view (scale bar, SO nm). **(E)** Segmentation-based quantification of atonosome volume in two yeast species, reflecting size."**" *p* value < 0.0 ; white horizontal bar, median; black vertical bar, IQR or middle 50%. **(F)** Eisosomes are observed at the PM and display a characteristic canoe-shaped morphology. **(G)** Spinning-disk confocal microscopy of C. *perkinsii* with the plasma membrane stained using FM4-64, under mock conditions (left panel) or following Palme treatment (4 µM, 10 min). Maximum-intensity projections along the z-axis. Scale bar, 5 µm. Scale bars, 100 nm unless otherwise indicated.

We then investigated whether this conservation would extend to other eukaryotes exhibiting complex life cycles and harboring a CW. To this end, we examined the membrane response to PalmC in the ichthyosporean *Chromosphaera perkinsii* ^43^. Ichthyosporeans are deep-branching opisthokonts that lie between animals and fungi and exhibit a mixture of fungal-like traits and transient multicellular stages reminiscent of early animal development ^44^. *C. perkinsii* is particularly notable because it is enclosed by a CW yet undergoes a palintomic developmental program with cleavage divisions, symmetry breaking and multicellular colony formation, features that show striking parallels with early animal embryogenesis ^45^. Remarkably, atonosomes could be visualized in *C. perkinsii* using FM4-64 PM staining following PalmC treatment **(Fig. 6G).** Their formation conditions and kinetics were notably similar to those observed in yeast **(Fig. S5G, Movie S12).** Altogether, the remarkable similarities in structure, formation conditions, and dynamics across Opisthokonts strongly suggest that atonosomes represent a conserved adaptive response to membrane stress.

## Discussion

Membrane clustering under altered PM homeostasis has long been described in the literature using diverse and seemingly unrelated terminology. Our unprecedented high-resolution ultrastructural analysis provides a framework for unifying past and present observations, demonstrating that atonosomes are PM-derived compartments with a defined lumen, organelle-like architecture, and dynamic responsiveness to membrane stress, arising from acute loss of membrane tension and from chronic dysregulation in PM homeostasis. Atonosomes induced by acute loss of membrane tension, that we termed as stress-induced, were first described as PM invaginations, PI(4,5)P_2_-enriched structures (PES) or large eisosome-derived structures (LEDS) ^12–14^. Our findings refine these previous nomenclatures, as atonosomes are detected not only with PI(4,5)P₂ probes but also with PS, PI4P and sterol markers, while eisosomes are dispensable for their formation.

The rapid formation of stress-induced atonosomes, their ATP-independent emergence, and the lack of mutants severely defective in their formation suggest that no dedicated active machinery is required and that biophysical changes at the PM are sufficient to initiate the process. Consistent with this, hyperosmotic shock has been shown to induce spherical invaginations in giant unilamellar vesicles *in vitro* ^46^. We therefore propose that abrupt changes in PM biophysical properties drive atonosome formation, which would act as a structural buffer for excess membrane generated by loss of PM tension.

The formation of stress-induced atonosomes coincides with TORC2 recruitment and inactivation, along with relocation of Slm1 and septins. We hypothesize that their recruitment is driven by fast changes in membrane curvature and/or lipid accessibility during atonosome formation, thereby creating a local hub for protein assembly ^47–50^. Supporting this, these proteins are spatially segregated within distinct regions of atonosomes: the base likely exhibits negative curvature, whereas other regions display null or positive curvature. Moreover, it was shown that septins directly bind PI(4,5)P_2_ and that PI(4,5)P2-containing liposomes can induce septin ring formation *in vitro* ^51,52^, also supporting the idea that lipid availability may enhance protein recruitment.

Although Slm1/2 have been implicated as TORC2 activators in previous studies, the molecular mechanism governing this regulation remains obscure. At first glance, the colocalization of Slm1 and TORC2 at stress-induced atonosome sites might seem to contradict the inactive state of TORC2. However, our findings reveal that TORC2 is distributed along the atonosome membrane, whereas Slm1 is confined to the base. This spatial segregation is likely sufficient to maintain TORC2 in an inactive state. By which mechanism TORC2 gets inactivated at atonosome sites, and the physiological significance of Slm1 and septins recruitment remain to be elucidated.

The fate/resolution of atonosomes remains unclear; we envision two non-mutually exclusive scenarios: either reabsorption of the compartment into the PM, resulting in flattening of the structure with its lumen becoming trapped between CW layers, or inward movement of the compartment into the cytoplasm followed by fusion of all or part with the vacuole for subsequent degradation. Interestingly, cryo-FIB–ET revealed images consistent with both scenarios: flattened structures with reduced lumen size and, free atonosomes that appear fully detached from the PM **(Fig. S1A)**. When investigating the fate of atonosomes in living cells, we found that stress-induced atonosomes disappeared within 15–20 minutes, whereas a subset of PalmC-induced atonosomes persisted for several hours. Because PalmC intercalates into the PM and generates an excess of lipidic material, cells may be challenged to resolve these oversized compartments. Several mechanisms could nevertheless contribute to atonosome resorption. Following hyperosmotic shock, activation of the HOG pathway stimulates glycerol production, restoring PM tension^53^. The resulting membrane stretching could promote the flattening of atonosomes at the PM. In contrast, for PalmC-induced atonosomes, we observed PS-positive membrane material detaching from these structures and subsequently accumulating at the vacuole, consistent with a possible trafficking route. However, better live imaging approaches and/or atonosome markers would be necessary to determine whether this represents motion of entire atonosomes, partial membrane fragments, or only lipids redistributed through membrane trafficking pathways.

Atonosomes also exist at steady state in eisosome mutants. These constitutive atonosomes share the same architecture as stress-induced ones and correspond to the previously described “dimple-like structures”, “cell wall invaginations” or “eisosome remnants” observed in *pil1Δ* cells ^9,15,16^. They were termed eisosome remnants mainly because some eisosomal proteins such as Sur7, Lsp1 and Slm1 were enriched at those sites ^9,54^. Here, we challenge this view: rather than representing residual eisosomes, we propose that eisosome sites no longer exist in the absence of the main scaffolding protein Pil1. Instead, the remaining eisosomal proteins would relocate to highly curved membrane structures: the atonosomes.

Eisosome mutants fail to keep TORC2 inactivated upon acute loss of PM tension, and deletion of *SLM1/2* in the *pil1Δ 6tspΔ* mutant abolishes the constitutive atonosome phenotype. We hypothesize that the loss of Pil1 disrupts Slm1 sequestration at eisosomes, resulting in erratic TORC2 activation and dysregulation of a major PM homeostasis mechanism. In support of this model, both constitutive activation of the TORC2 substrate Ypk1 and overexpression of *SLM1* induce persistent atonosome formation ^55^ likely through disrupted lipid biosynthesis, endocytosis, and vesicular trafficking ^11,12^. We performed a confocal screen for PM architecture defects across a deletion library and revealed 30 hits presenting lipid clustering at the PM using PI(4,5)P_2_ and PS reporters. Consistent with the idea that constitutive atonosomes results from lipid homeostasis imbalance, many of the genes identified in our genetic screen encode proteins involved in lipid metabolism, such as *ORM2, SAC1* and *ERG2.* Loss of *ORM2* or *SAC1* may result in an excess of sphingolipids or phosphoinositides, whereas deletion of *ERG2* would lead to the accumulation of sterol intermediates. Altogether, we propose that changes in the lipidic landscape of the PM, caused by TORC2 hyperactivity and/or excess of lipids, would chronically lower membrane tension, thereby mimicking acute stress conditions and stabilizing the atonosome phenotype.

Beyond lipid regulation mutants showing constitutive atonosomes, we also identified eight genes involved in vesicular transport and endocytosis and three encoding vacuolar H^+^-ATPase (V-ATPase) subunits or assembly factors **(Table 1).** Because vesicular trafficking is essential for lipid turnover through continuous endocytic cycling, its disruption is expected to perturb membrane homeostasis, thus promoting atonosome formation ^56^. Furthermore, the vacuole functions as a key osmotic buffer in yeast and is the end point of the endocytosis pathway ^57,58^. A non-functional V-ATPase at the vacuole surface will shift the osmotic load directly onto the PM which may result in chronic stress and atonosome formation. Interestingly, it was also recently proposed that loss of V-ATPase activity impairs inositol metabolism in yeast ^59^, further confirming that our screen reflects broad lipid homeostasis defects. Collectively, this screen provides an interesting framework to study how diverse cellular pathways impact membrane organization.

Atonosomes appear to represent a conserved response to acute PM tension decrease in *S. cerevisiae, S. pombe and C. perkinsii* despite their considerable phylogenetic distance. Moreover, PM macroinvaginations described in filamentous fungi – which encompass a broad diversity of structures and have been proposed to function as a reservoir for hyphal growth - may constitute another manifestation of atonosomes in cell wall-enclosed organisms ^60^. Strikingly, atonosome-like structures of unknown function have also been recently observed in plant cells by cryo-FIB-ET, highlighting a potential cross kingdom biological significance ^61^. The presence of a CW and its requirement for proper atonosome organization represents a key finding: this distinguishes atonosomes from simple membrane invaginations and establishes them as bona fide architectural entities. The high plasticity of the CW along the atonosome membranes suggests that a substantial part of it is not fully polymerized and available for dynamic changes. This paves the way for future investigations of membrane dynamics that account for the CW plastic properties. Interestingly, atonosomes protrude when the CW is absent, recalling recently discovered mammalian blebbisomes and zombosomes, involved in cell migration and cancer development ^62,63^. Overall, atonosomes are the first discovered internal compartment enclosed with CW material and the fact that they form largely based on membrane biophysics suggest potential pro-endocytosis behavior as well as shed light on origins of organelle generation, pointing to a largely unexplored layer of cellular biology.

## Material and Methods

### Cell culture and treatments

Yeast strains were generated using standard homologous recombination techniques. *S. cerevisiae* strains used in this study were derived from the BY4741 or TB50 background. Cells were routinely cultured in synthetic complete (SC) medium buffered to pH 6.2 with 0.1 M Sorensen buffer and supplemented with 2% glucose and the appropriate amino acids, with shaking at 30 °C. All experiments were performed using logarithmically growing S*. cerevisiae* cells (OD₆₀₀ = 0.6–1).

*S. pombe* cells were cultured at 30 °C with shaking in Edinburgh minimal medium supplemented with amino acids (EMM-ALU) (0.67% Difco yeast nitrogen base without amino acids, 2% glucose, and amino acids supplemented as required)^64^. All experiments were performed using logarithmically growing *S. pombe* cells (OD₆₀₀ = 0.6–0.8).

*C. perkinsii* cells were cultured at 26 °C in liquid medium (yeast extract 3 g/l, malt extract 3 g/l, peptone 5 g/l, glucose 10 g/l, NaCl 20 g/l) in a rectangular canted neck cell culture flask with vented cap (Falcon; 353108), protected from light, and refreshed weekly (1/10 dilution).

For hyperosmotic shock experiments, cultures were diluted with medium containing 2 M sorbitol to achieve a final concentration of 1 M sorbitol. Alternatively, for cryo-FIB-ET experiments, glycerol was added directly to the culture medium to a final concentration of 5% (0.5 M). For PalmC treatment, PalmC was added to the culture medium to a final concentration of 8 µM.

### Preparation of uniform cell layer for cryo-FIBM/SEM

Four µL of *S. cerevisiae* or *S. pombe* cells at OD₆₀₀ = 0.8-1 were deposited onto glow-discharged holey carbon-coated Quantifoil R 2/2, 200-mesh copper grids. Cells were vitrified using a GP2 plunge freezer (Leica Microsystems). Back blotting was performed using Grade 595 filter paper. Blotting conditions were optimized to facilitate subsequent cryo FIB milling : 22 °C, 60–70% humidity, a wait time of 10 s, and a blot time of 2–3 s. Grids were then rapidly vitrified in liquid ethane. Subsequently, grids were clipped into cryoFIB autogrids (Thermo Fisher Scientific, TFS). This procedure yielded well-preserved grids uniformly covered with a single layer of yeast cells **(Fig. S6Aiii)**. Grid bars remained visible, and all grid squares were suitable for lamella preparation **(Fig. S6Ai-iv).**

### Cryo-FIBM/SEM workflow

Specimen thinning has been revolutionized over the past fifteen years by cryogenic focused ion beam milling combined with scanning electron microscopy (cryo-FIB-SEM) ^65,66^.

Cryo-FIB milling was performed using an Aquilos 2 dual beam instrument (TFS) at the Electron Microscopy Facility (EMF), University of Lausanne (UNIL), Switzerland. Inside the FIB chamber, grids were first sputter-coated with platinum for 60 s, 30 mA, followed by deposition of an organometallic platinum layer using the integrated gas injection system (GIS) for 1 min 30 s, and a second 60 s platinum sputter coating. This extensive coating procedure was performed to protect the specimen surface, reduce charging during milling, and minimize curtaining artifacts.

At a stage tilt of 17°, grid squares containing a uniform and single layer of cells were positioned at the coincidence point for serial FIB slicing and SEM imaging. Lamellae of 150–200 nm thickness (10–20 µm width and 10–20 µm depth) were programmed using AutoTEM software (TFS). Rectangular milling patterns were applied with beam currents ranging from 1 nA to 300 pA for rough milling, 100 pA for fine milling, and 30 pA for final polishing **(Fig. S6Aiv)**. Milling was automatically performed sequentially at each site, followed by automated overnight polishing. Up to 50 lamellae could be generated automatically in 24h. Stress-relief trenches were occasionally milled when the cell layer covered the entire grid square surrounding the lamella position **(Fig. S6Aiv)**. Based on our experience, approximately 75% of lamellae reached the target thickness of 150–200 nm and were subsequently suitable for imaging **(Fig. S6Av-vi)**.

### Cryo-electron tomography

Grids containing the lamellae were manually oriented perpendicular to the tilt axis prior to loading into the transmission electron microscope. Tilt series were acquired on a 300 kV Titan Krios E-CFEG (TFS) equipped with a Selectris X post-column energy filter (TFS) and a Falcon 4i direct electron detector (TFS) at the Dubochet Center for Imaging (DCI), Lausanne, Switzerland. Samples were tilted from −50° to +50° in 2° increments according to the dose-symmetric Hagen scheme ^67^. The 0° tilt angle was defined by accounting for the intrinsic tilt of the lamellae resulting from the milling angle. Tilt series were acquired using Tomo5 (TFS) at nominal magnifications of 42,000× or 64,000×, corresponding to pixel sizes of 3.1 Å and 2.0 Å, respectively, with a cumulative electron dose of approximately 120-150 e⁻/Å². Tomograms were reconstructed live using TomoLive (TFS) or the Scipion pipeline for initial quality assessment.

### Tomogram processing and segmentation

Frame alignment and dose weighting of tilt series were performed using MotionCor2 v1.6.4 according to the cumulative electron dose ^68^. Tilt series alignment was carried out using the patch-tracking method implemented in etomo, IMOD v4.11.12 ^69^. Tomograms were reconstructed in IMOD v4.11.12 using weighted back projection, and SIRT-like filtering was applied for visualization purposes.

Automated segmentation was performed using a conventional neural network algorithm implemented in EMAN2 v2.99.66 ^70^. Training datasets were generated for ribosomes and the cell wall. For membrane segmentation, tomograms containing atonosomes were automatically processed using MemBrain ^71,72^.

For visualization purposes, endoplasmic reticulum membranes and the inner membrane of the atonosomes were manually segmented using the Drawing and Interpolator tools in IMOD v4.11.12. Isosurface renderings of cellular features were generated using UCSF ChimeraX v1.6.1 ^73^. Small particles corresponding to false-positive segmentations were removed using the *Hide Dust* tool in ChimeraX v1.6.1.

### CryoCLEM

Clipped grids were mounted on a cryo cassette for the Leica Thunder cryo-LM (Leica Microsystems) and transferred to the cryostage for imaging. A complete overview map (11 x 12 tiles) of the grid was acquired using bright field and reflected light with a 50x objective (NA 0.75, pixel size 260 nm), using many focus points set to ensure sharp focus across the entire surface. Fluorescence z-stacks of the mCh-2xPH signal were collected at the center of the grid. Fluorescence data was imported into Maps 3.29 (TFS) in the Aquilos2 setup and aligned to the ion beam image using three reference points (milling bevel, holes in the grids). The overlay enabled direct stage navigation to regions of interest. The fluorescence/ion correlated images were then uploaded into Maps in the Titan Krios setup and further correlated to the TEM Atlas, giving a broad overview of the fluorescence signal matching lamellae positions. Post acquisition, precise positioning of the fluorescence signal onto the Search maps obtained by TEM was done using Napari v0.5.4^74^.

### High-pressure freezing, freeze substitution and ultramicrotomy

*S. cerevisiae* cell pellets were dispensed on the 200-µm side of a 3-mm type A gold/copper platelet, covered with the flat side of a 3-mm type-B aluminum platelet (Leica Microsystems). The sample was vitrified by high-pressure freezing (HPF) using an ICE system (Leica Microsystems) in which cells were subjected to a pressure of 210 MPa at −196 °C.

Freeze substitution for conventional TEM imaging, was performed at the Electron Microscopy Facility of the Institute of Structural Biology, Grenoble, France. HPF samples were then gradually warmed to −60 °C at a rate of 2 °C/h using an AFS2 system (Leica Microsystems). After 8–12 h at −60 °C, the temperature was increased to −30 °C at 2 °C/h and subsequently raised to 0 °C within 1 h to enhance osmium fixation and improve membrane contrast. Samples were then cooled back to −30 °C within 30 min and rinsed four times in pure acetone. Samples were infiltrated with progressively increasing concentrations of resin (Embed-812, EMS) in acetone (1:2, 1:1, and 2:1 [v/v]) for 2 h at each ratio while gradually warming to 20 °C. Pure resin was subsequently added at 20 °C and polymerized at 60 °C for 48 h. 70 nm thick sections were prepared using a diamond knife (Diatom) and a UC7 ultramicrotome (Leica Microsystems) and collected on carbon-coated 200-mesh copper grids (Agar Scientific) at the DCI, Geneva, Switzerland. Sections were post-stained with 2% aqueous uranyl acetate for 5 min, rinsed in water, incubated in lead citrate for 5 min, and rinsed again. Imaging was performed using a Talos L120C (TFS) operated at 120 kV.

For serial sectioning, samples were freeze-substituted using an AFS2 system (Leica Microsystems) at the EMF, Lausanne, in 0.2% uranyl in acetone at −90 °C for 10 hours. Temperature was progressively raised to - 45°C over 14 hours. Samples were rinsed in pure acetone three times at -45°C. Samples were subsequently infiltrated with gradually increasing concentrations of Lowicryl HM20 resin (EMS, Hatfield, PA, US) in acetone (10%, 25%, and 75%) for 4 h at each concentration while gradually warming to −25 °C. Pure resin was then added at −25 °C for 20 h. Resin polymerization was performed under UV illumination for 48 h at −25 °C, followed by a gradual temperature increase to 20 °C over 9 h under continuous UV exposure. Final polymerization was completed at room temperature for 24 h. Serial sections were prepared using a UC7 ultramicrotome (Leica Microsystems). Ribbons of 7–8 sections were collected on copper slot grid 2x1mm (EMS, Hatfield, PA, US) coated with a PEI film (Sigma, St Louis, MO, US) and imaged using Tecnai 12 (TFS) operated at 120 kV. Regions of interest were identified in each section manually. Images of the same atonosome across serial sections were aligned using the Align Serial Section pipeline in etomo, IMOD v4.11.12. Further contrast adjustment was done in Fiji v1.54 ^75^

### Live-cell microscopy, treatments and image processing

For all fluorescent microscopy experiments, all imaging was performed with a Nikon Ti / CSU-W1 Spinning Disc Confocal Microscope with a 100x oil immersion objective, 3i software and sCMOS camera with 1200X1200 pixels (FOV 11x11um, pixel size 0.11µm).

*S. cerevisiae* or *S. pombe* cells were grown to mid-log phase in appropriate low fluorescent culture medium and deposited in Concanavalin A-coated VI 0.4 µ-Slides (Ibidi) washed with media at room temperature. Imaging was performed at 100% laser power at 561 nm excitation for mCherry and 488 nm for GFP, with a 150ms exposure. For live imaging of PalmC and hyperosmotic shocks, culture medium and cell excess are removed and imaging positions were rapidely selected proximal to the channel opening where the fresh medium is added. Imaging starts without treatment and medium containing 1M sorbitol or 8 µM PalmC is added to the chamber while imaging. For fast imaging, z stacks spanning 5-8 µm are taken every 2-8 seconds depending on the number of channels and slices in the stack. For long term imaging of atonosome fate, images are taken every minute.

To investigate the effect of hypoosmotic shock on atonosomes, two experimental setups were used: For atonosomes induced by hyperosmotic shock, structures were generated in ibidi chambers as described above. Hyperosmotic medium was then removed and replaced with standard culture medium to induce hypoosmotic conditions. For PalmC-induced atonosomes, cells were pre-adapted by growing them in culture medium containing 1 M sorbitol for several hours, resulting in a smooth plasma membrane. Cells were then transferred to an Ibidi chamber. Culture medium supplemented with 1M sorbitol and 8 µM PalmC was added. Once atonosomes were formed, hypoosmotic shock was applied by exchanging this medium for standard culture medium. For calcofluor white staining, 4 uM calcofluor white was added to the culture for 5 minutes with shaking 30 °C prior deposition in the in ibidi chamber. For TORC2 staining (Avo2-Halo and Avo3-Halo) Janelia halo ligands combined to alexa fluor 646 (Promega) was added to the culture at 200 nM for 1 to 2 hours, at 30 °C with shaking. Cells were then washed twice in culture medium prior deposition in ibidi chamber.

For *C.perkinsii,* samples were prepared from a dense asynchronous culture and diluted 10x in culture medium. Colonies shown in figures are all at the unicellular stage. Cells were stained with 2µM FM4-64 Dye (Thermofischer; T13320) reconstituted 3h before imaging in Milli-Q water, and imaged directly after staining. PalmC was added at 4µM (in DMSO) in the solution during imaging. Cells were mounted in a 4-well µ-Dish, 35 mm Quad Uncoated (Ibidi; 80411) at 26 °C. Imaging was performed at 100% laser power at 561 nm excitation, with a 100 ms exposure. Images were acquired in 3D with 0.5 µm z- spacing for a total range of 20µm in depth and movies were acquired with a 30 sec interval for 30 min.

All images were contrast-adjusted and zoomed for presentation purposes only. Additionally, in case of bleaching, the contrast was adjusted overtime with the “bleach correction* plugging using histogram matching parameter ^76^. For long term imaging (1h or more) the potential drift in focus was corrected using Fast4Reg plugin ^77,78^ in Fiji.

### Yeast protoplasting

Cells were grown to OD 0.8-1 in SC medium, pelleted at 1500g for 1’30 min and resuspended in SC with 1M sorbitol. Cell wall was removed using 0.1 mg/mL zymolyase, 10% DTT, incubated at 30 °C for 45 min. Cells were washed with SC 1M sorbitol twice and imaged in ibidi chambers similarly to intact cells. PalmC treatment was performed at 8 µM in SC 1M sorbitol for 5 min.

### WB sample preparation and detection of phosphoproteins on WB

Yeast culture aliquots were processed according to standard TCA-urea extraction procedures. In brief, culture aliquots were incubated with 5% TCA on ice for at least 10 min, before pelleting and drying cells with ice-cold acetone. Cells were lysed by bead beating in a urea buffer (25 mM Tris pH 6.8, 6 M Urea, 1% SDS), and boiled for 5 min. Denatured lysates were mixed with 2x sample buffer (125 mM TRIS pH 6.8, 20 vol% glycerol, 2% SDS, 0.02% Bromphenol Blue, 200 mM DTT), and boiled again before analyzing them via WB. Mammalian cells were lysed with a lysis buffer (40 mM HEPES, 10 mM Na-PPi, 10 mM Na-β-glycerophosphate, 4 mM EDTA 1% Triton X-100, pH 7.4) supplemented with 1× Halt Protease & Phosphatase Inhibitor Cocktail (Thermo Fisher Scientific) and kept at −20 °C until further analysis. A Pierce BCA Protein Assay (Thermo Fisher Scientific) was used to determine protein concentration in the samples. In all, 15–50 µg of total protein of each sample was used for further analysis. The 5x sample buffer (312.5 mM Tris, 10% SDS, 50% glycerol, 25% β-mercaptoethanol, 0.1% bromophenol blue, pH 6.8) was added, and samples were denatured at 95 °C for 5 min. Protein lysates were separated on 10% SDS-page gels and blotted onto nitrocellulose membranes using the iBlot2 Gel Transfer System (TFS). Membranes were blocked with BSA, incubated with primary antibodies overnight at 4 °C, washed, and incubated with secondary antibodies for 1 h at room temperature, using TBS based buffers. Membranes were developed using an Odyssey imaging system (LI-COR), and signal intensities were quantified using Fiji. Calculations were performed in Microsoft Excel, and data were plotted using GraphPad Prism (10.2.3).

### Segmentation-based measurement of atonosomes volume

Segmentation masks obtained with MemBrain were processed using the MorpholibJ plugin (v1.6.5)^79^ in Fiji. First, connected components were identified in 2D, and the components corresponding to the atonosomes were manually selected and merged. Several morphological parameters were measured in 3D, including minimum and maximum coordinates along the x, y, and z axes, volume, surface area, and mean breadth, all initially expressed in pixels. These measurements were subsequently normalized according to the pixel size.

As the atonosomes are larger than the lamella thickness, volume estimation accounted for an average lamella thickness of 200 nm. The volume was estimated as:

*Volume = (Max X − Min X) × (Max Y − Min Y) × 200 nm*

An extrapolated volume was also calculated assuming that atonosomes are spherical:

*Extrapolated volume = 4/3 πr³,* where *r* is the average radius.

Mean breadth, also referred to as mean width or mean diameter, is a morphological feature that quantifies object diameter, taking into account concavities. Therefore, this metric also reflects structural complexity.

### Library creation via synthetic genetic array and HTP imaging

The yeast Knockout library and DAMP essential knockdown libraries ^26^ were arranged in 1536 spot arrays on YPD-G418 plates, and then mated with the query strain yMS2085 (MatAlpha his3Δ1 leu2Δ0 met15Δ0 ura3Δ0 LYS2+ can1Δ::GAL1pr-SceI::STE2prSpHIS5 lyp1Δ::STE3pr-LEU2) ^80^ containing mCherry-tagged tracker of PI(4,5)P_2_ and GFP-tagged tracker for PS. The library creation then followed the standard synthetic genetic array protocol ^81^ with LEU selection for the Alpha mating type. The final library was transferred to a liquid culture in 384-well plates with 35 µL final selection media. After three days of growth, 15 µL glycerol were added to a final concentration of 15% and cells were stored at-80 °C. Prior to imaging, cells were transferred to polypropylene 384-well plates (Greiner™) containing 35 µL of complete synthetic medium, and grown overnight. From this culture, 1 µL was transferred to 35 µL fresh complete synthetic medium in optical bottom 384-well plates (Revity) and cells were grown for 6 h at 30 °C without shaking prior to imaging. One round of imaging (approximately 45 minutes, 6 images per well) was done without any additional treatments to observe effects of gene deletion/down-regulations on probe localization, and another two rounds were done after addition of 8mM of PalmC. Imaging was performed on an Olympus IX83 microscope coupled to a Yokogawa CSU -W1 spinning -disc confocal scanner with dual prime - BSI sCMOS cameras (Photometrix). The 16-bit images were acquired with three illumination schemes for brightfield, green fluorescence, and for red fluorescence. For green fluorescence, we excited the sample with a 488 nm laser (Coherent 150 mW) and collected light through a bandpass emission (Em) filter (525/50 nm, Chroma ET/m). For red fluorescence, we excited at 561 nm (Coherent 150 mW) and used a bandpass emission filter (609/54 nm, Chroma ET/m). One multiband dichroic mirror was used for the three illuminations. Imaging was performed with a ×60/1.42 numerical aperture (NA), oil -immersion objective (UPLSAPO60XO, Olympus). Following acquisition, cells were manually curated for membrane defects or segmented and their fluorescent signal (median, average, minimum, maximum, 10th, 20th, …, 90th percentile fluorescence) were identified using previously reported scripts in ImageJ ^82^.

### Multiple protein sequence alignment

Multiple sequence alignment and ortholog findings was performed with the Orthologous Matrix (OMA) server (https://omabrowser.org/) (Zahn-Zabal et al., 2020). Results were analyzed and visualized with Jalview (v2.11.5.1, Waterhouse et al., 2009)

### Statistics and reproducibility

Cryo-FIB/ET and TEM on resin sections were performed independently at least twice for each strain or treatment, yielding consistent conclusions.

Fluorescence microscopy observations were performed at least 3 independent times for each strain and treatment. The numbers of atonosomes present in the cells were manually measured over 65 cells and 3 different independent experiments.

Western blot results for TORC2 activity (pT662 Ypk1) is presented as mean values and SD from at least three independent replicates (or at least two replicates for supplementary figures).

Statistical analysis was performed using the student’s *t-test*.

## Acknowledgment

We thank all members of the Loewith and Boland laboratories for their support, insightful discussions, advice, and encouragement throughout this project. We are also grateful to the Levy, Martin, and Dudin laboratories for their great contributions to scientific discussions and for participating in interdepartmental collaborations.

We thank the staff of the microscopy platforms, without whom this project would not have been possible. In particular, we acknowledge the Electron Microscopy Facility (EMF) Lausanne, with special thanks to C. Genoud for her continuous support throughout the project. We also thank the Dubochet Center for Imaging (DCI) Lausanne, especially B. Beckert, E. Uchikawa, I. Mohammed, and S. Nazarov, for their expertise and assistance. We thank the DCI Geneva team for their support with sample preparation and scientific advice, with special thanks to A. Howe, S. Barrass, and C. Bauer. We further thank J. Bosset from the Photonics Facility of the University of Geneva (UNIGE) for technical support. We thank the computing department of the UNIGE for providing an infrastructure to perform cryo-EM analysis; N. Roggli for maintaining the computing infrastructure in the Molecular and Cellular Biology department. This project was made possible through the support of the Bryn Turner-Samuels Foundation, in partnership with the Foundation for Research in Biology and Medicine. We acknowledge support from EMBO for E. Bauda postdoctoral fellowship (ALTF 245-2024).

This work made use of the Cellular Electron Microscopy platform at the Grenoble Instruct-ERIC Center (ISBG; UAR 3518 CNRS–CEA–UGA–EMBL), within the Grenoble Partnership for Structural Biology. We particularly thank C. Moriscot, B. Gallet, and G. Schoehn for their assistance and expertise.

## Authors’ Contribution

Conceptualization: E.B, R.L, A.B. Methodology Validation: E.B, R.L, A.B, A.A, M.T, S.M, O.D, E.L. Investigation: E.B, A.A, M.T, J.CS, A.G, N.S, M.G, P.L, C.G, G.S, J.D. Resources: R.L, A.B, J.D. Data curation: E.B. Writing—original draft: E.B. Writing—review and editing: R.L, A.B, E.B, A.A, M.T, N.S, J.CS, S.M, O.D. Project administration: R.L, A.B, E.B Funding acquisition: R.L, A.B, E.B

---

**Figure S1.**
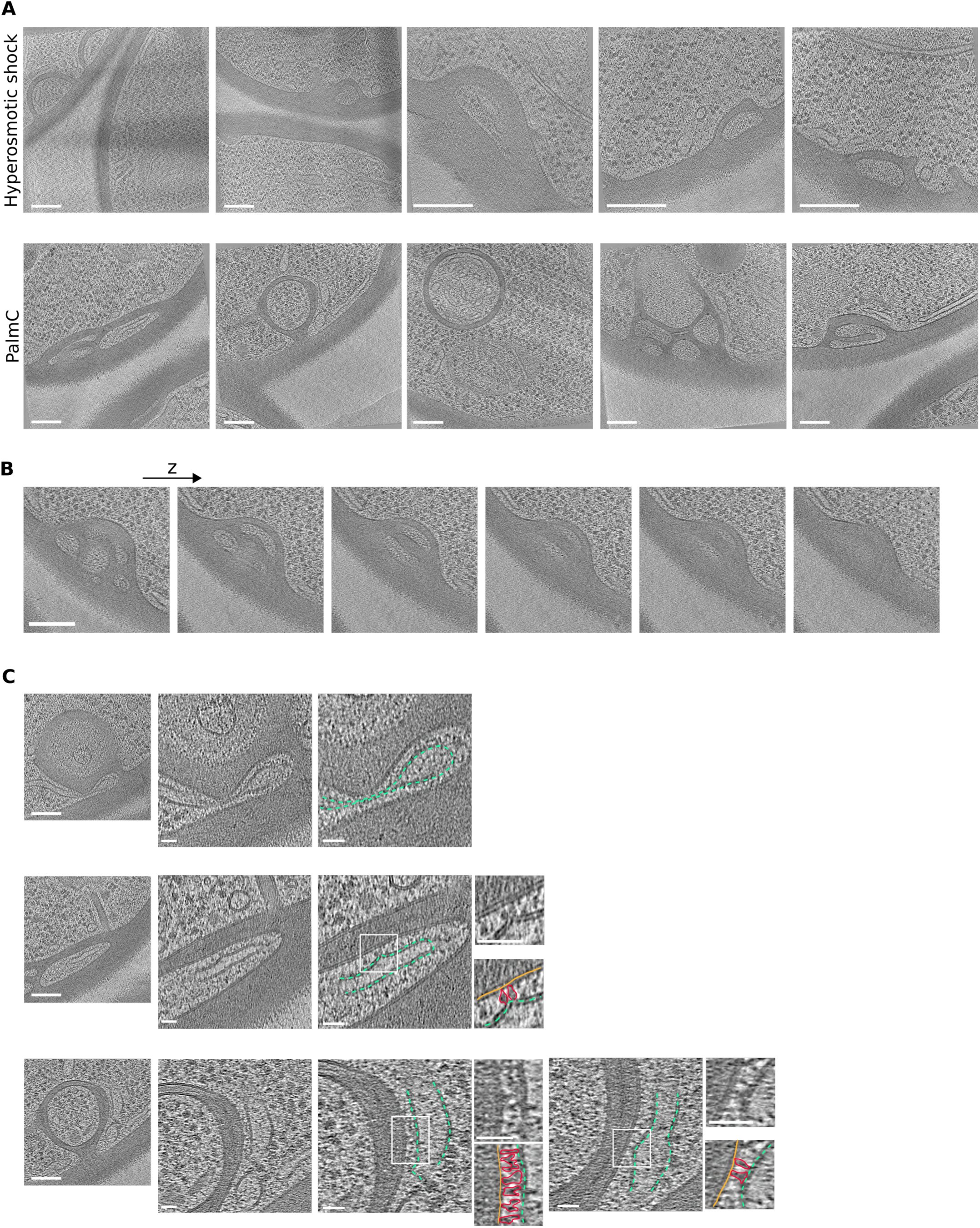
In situ electron micrographs showing diversity of morphology of *S. cerevisiae* atonosomes. **(A)** Cryo-tomographic slices of atonosomes formed under hyperosmotic shock or Palme treatment. **(B)** Cryo-tomographic slices of an atonosome with increment in the z axis, corresponding to lamella thickness. Cell wall appears to fully enclosed the compartment on one side. **(C)** Cryo-tomographic slices showing the close associations of ER (dashed green lines) and atonosomes membranes (yellow lines). Protein densities appear to connect the two structures (red structures). Magnified view scale bars, 25 nm. Scale bars, 100 nm unless otherwise indicated.

**Figure S2.**
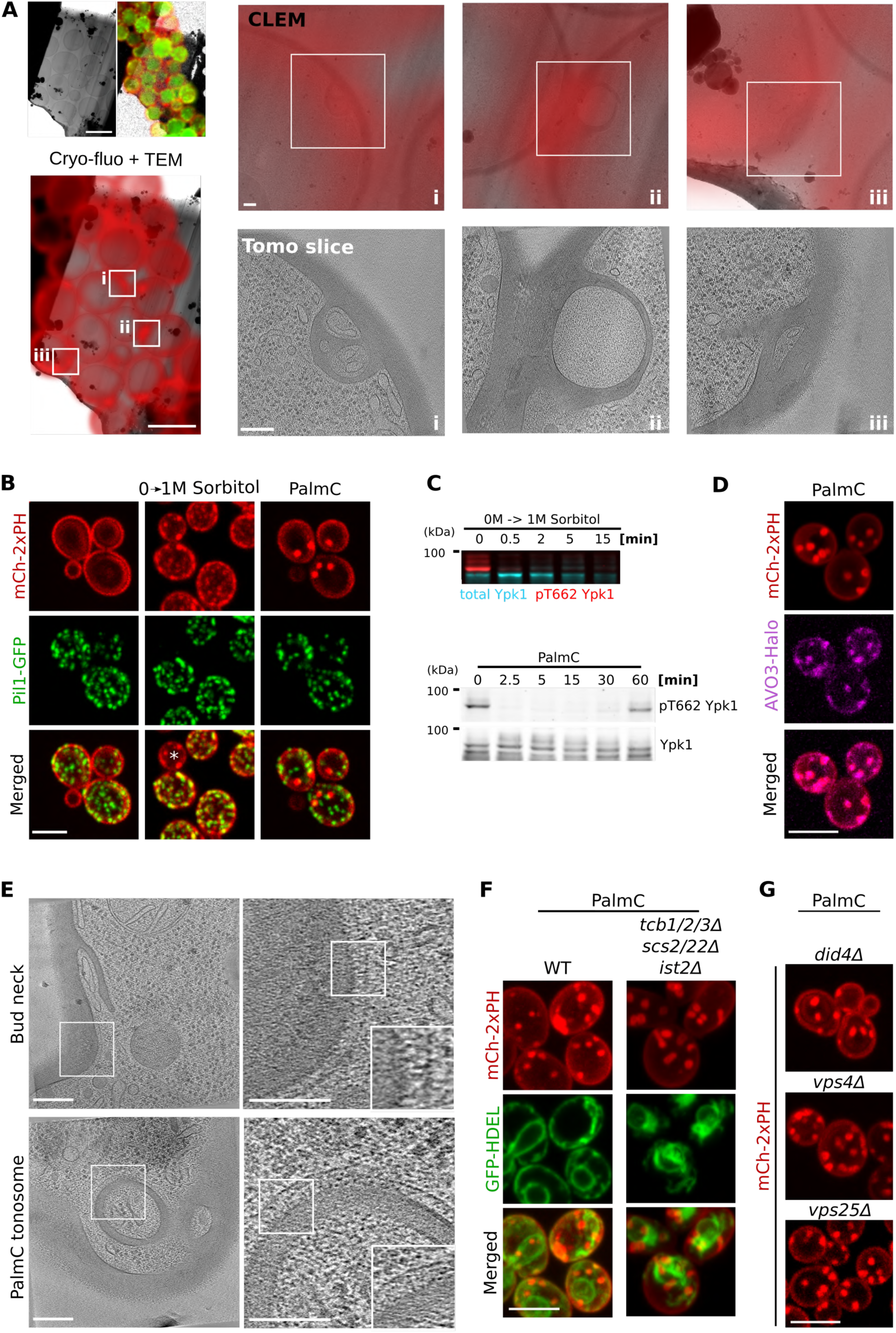
Atonosomes are eisosome-independent structures that form without active machinery and correlate with TORC2 inactivation. **(A)** CLEM correlates PM fluorescence puncta with atonosomes in cryo-lamellae of *WT* cells expressing the Pl(4,5)P_2_ reporter (mCh-2xPH). Upper left panels: TEM image of a cryo-FIB lamella with corresponding cryo-wide-field fluorescence (PM, mCh-2xPH; Pil1-GFP). Lower right panel: CLEM overlay showing three regions (i-iii) where PM puncta coincide with atonosomes detected by cryo-ET. Scale bar, 5 µm, magnified views, 100 nm. **(B)** Spinning-disk confocal microscopy of cells co-expressing endogenously Pil1-GFP and the Pl(4,5)P_2_ reporter (mCh-2xPH) under hyperosmotic shock (1M) sorbitol for 1 min) or Palme (8 µM for 5 min). White star shows a bud. **(C)** Western blot analysis showing TORC2 downregulation following hyperosmotic shock or Palme treatment. TORC2 activity was assessed by relative Ypk1 phosphorylation levels (pT662 Ypk1). **(D)** Spinning-disk confocal microscopy of cells expressing a Pl(4,5)P_2_ reporter (mCh-2xPH) from a plasmid and an endogenous Avo3-Halo (TORC2 subunit) lapebed with a Halo fluorescent ligand (646), under Palme treatment (8 µM for 5 min). **(E)** Cryo-tomographic slices showing septin alignment at the bud neck and along the Palme-induced atonosome outer membrane. Insets show magnified views of septin ring units. Scale bars, 100 nm; insets, 50 nm. **(F)** Spinning-disk confocal microscopy of Palme treated cells (8 µM for 5 min) expressing endogenous GFP-HDEL, targeting ER, and a Pl(4,5)P_2_ reporter (mCh-2xPH) in a WT background or in absence of six majors ER-PM tether proteins. **(G)** Spinning-disk confocal microscopy of indicated ESCRT deletion mutants expressing an endogenous Pl(4,5)P_2_ reporter (mCh-2xPH) under Palme treatment (8 µM for 5 min). All fluorescence images are maximum-intensity projections along the z-axis. Scale bars, 5 µm unless otherwise indicated.

**Figure S3.**
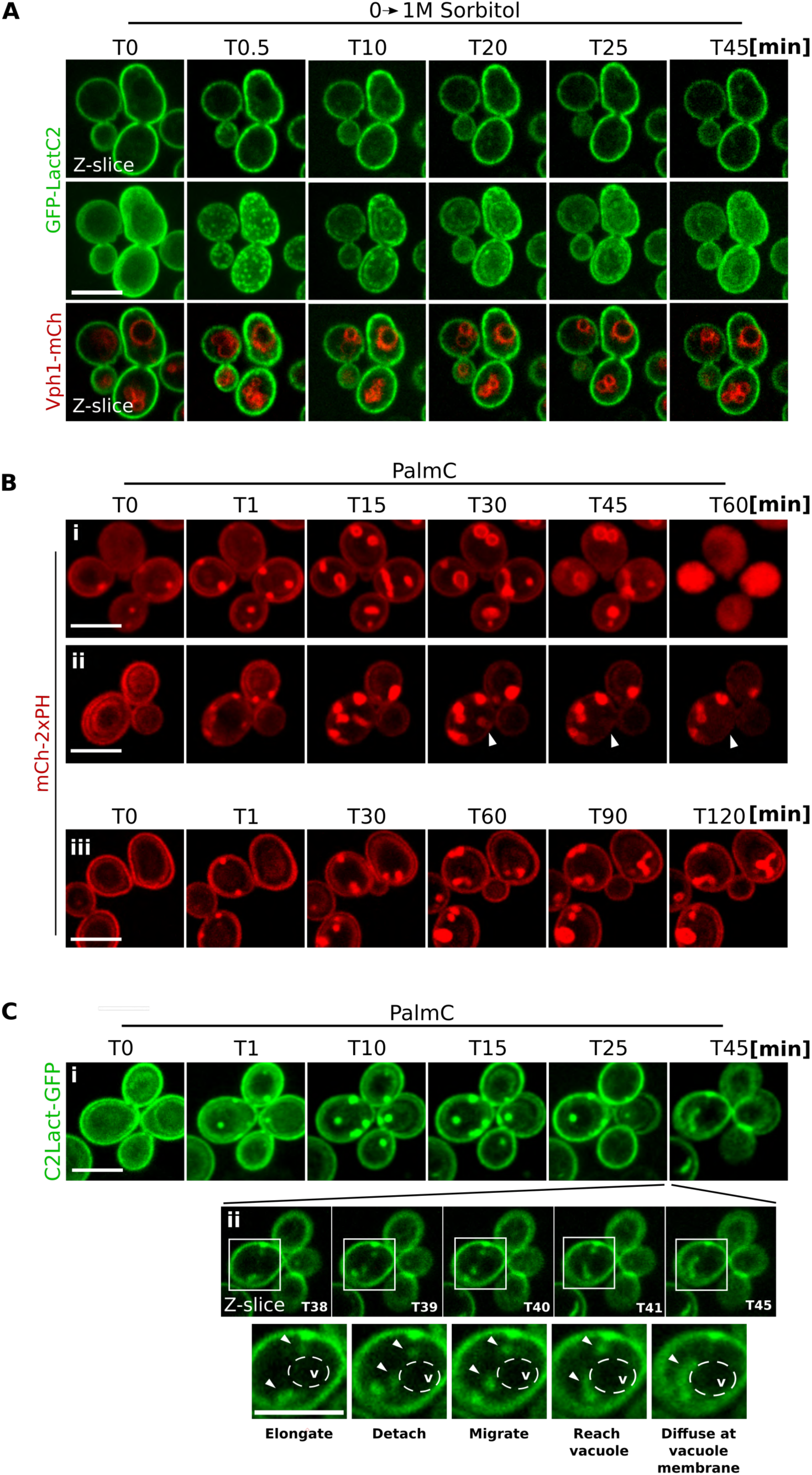
Fate of stress-induced atonosomes. **(A)** Live spinning-disk confocal microscopy showing the fate of atonosomes induced by hyperosomotic shock in WT cells expressing a PS reporter (GFP-LactC2) on a plasmid and Vph1-meherry at the vacuole endogenously. **(B)** Long-term live spinning-disk confocal imaging of *WT* cells expressing the endogenous Pl(4,5)P_2_ reporter (meh-2xPH) following Palme treatment. Over time, the Pl(4,5)P_2_ signal occasionally collapses into the cytoplasm **(i),** some atonosomes disappear (white arrows) **(ii),** whereas others remain stable for more than 2 hours **(iii)**. **(C)** Long-term live spinning-disk confocal imaging of *WT* cells expressing a PS reporter (GFP-Lacte2) on a plasmid following Palme treatment. Upon atonosome disappearance, GFP-Lacte2 signal appears on a structure resembling the vacuole **(i).** Tracking of GFP-LactC2-positive puncta reveals their movement from atonosomes to the apparent vacuolar surface (dashed circle, *v)* **(ii)**. Unless otherwise indicated, all fluorescence images are maximum-intensity projections along the z-axis. Scale bars, 5 µm.

**Figure S4.**
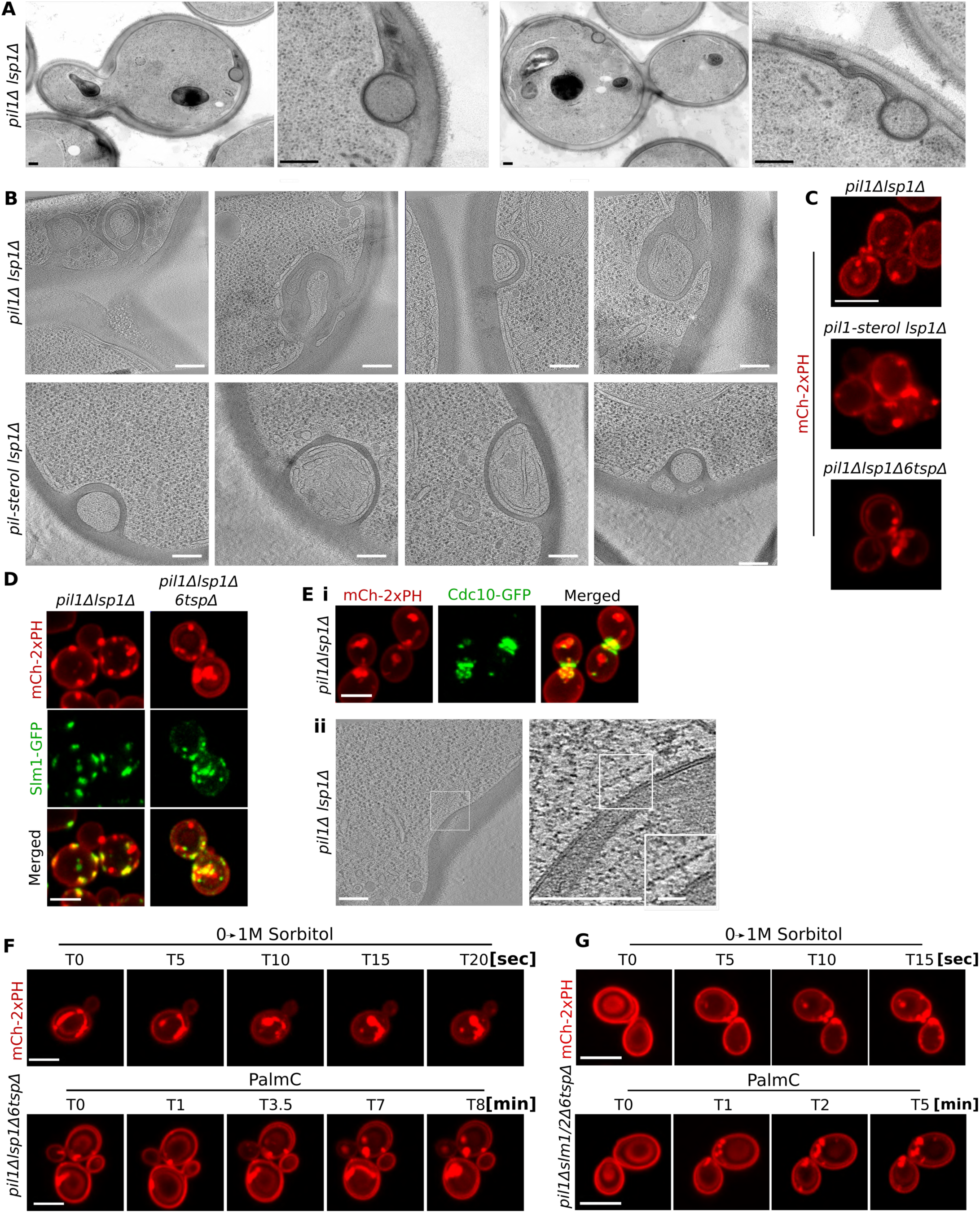
High-resolution imaging and stress-dependent dynamics of constitutive atonosomes. **(A)** TEM images of resin-embedded *pil1Δ. lsplΔ.* cell sections showing constitutive atonosomes in steady state. Scale bar, 200 nm. **(B)** Cryo-tomographic slice showing constitutive atonosomes in *pillΔ. lsp1Δ* and *pi/1-stero/ /splΔ.* cells. Scale bar, 100 nm. **(C)** Spinning-disk confocal microscopy of indicated eisosome mutants expressing a Pl(4,5)P_2_ reporter (mCh-2xPH) endogenously or From a plasmid. **(D)** Spinning-disk confocal microscopy of indicated eisosome mutants co-expressing Slm1-GFP and the Pl(4,5)P_2_ reporter (mCh-2xPH). **(E)(i)** Spinning-disk confocal microscopy of pil1n lsp1n cells co-expressing the septin Cdc10-GFP and a Pl(4,5)P_2_ reporter (mCh-2xPH) endogenously. (ii) Cryo-tomographic slices of septin-like alignment on a constitutive atonosome OM and magnified view of a septin unit (scale bar, 100 nm and 20 nm respectively). **(F)** Live spinning-disk confocal microscopy of *pil1Δ lsp1Δ6tspΔ* cells expressing a Pl(4,5)P_2_ reporter (mCh-2xPH) from a plasmid, subjected to hyperosmotic shock or Palme treatment. **(G)** Live spinning-disk confocal microscopy of *pillΔslm1/2Δ6tspΔ* expressing a Pl(4,5)P_2_ reporter (mCh-2xPH) from a plasmid, subjected to hyperosmotic shock or Palme treatment. All fluorescence images are maximum-intensity projections along the z-axis. Scale bars, 5 µm unless otherwise indicated.

**Figure S5.**
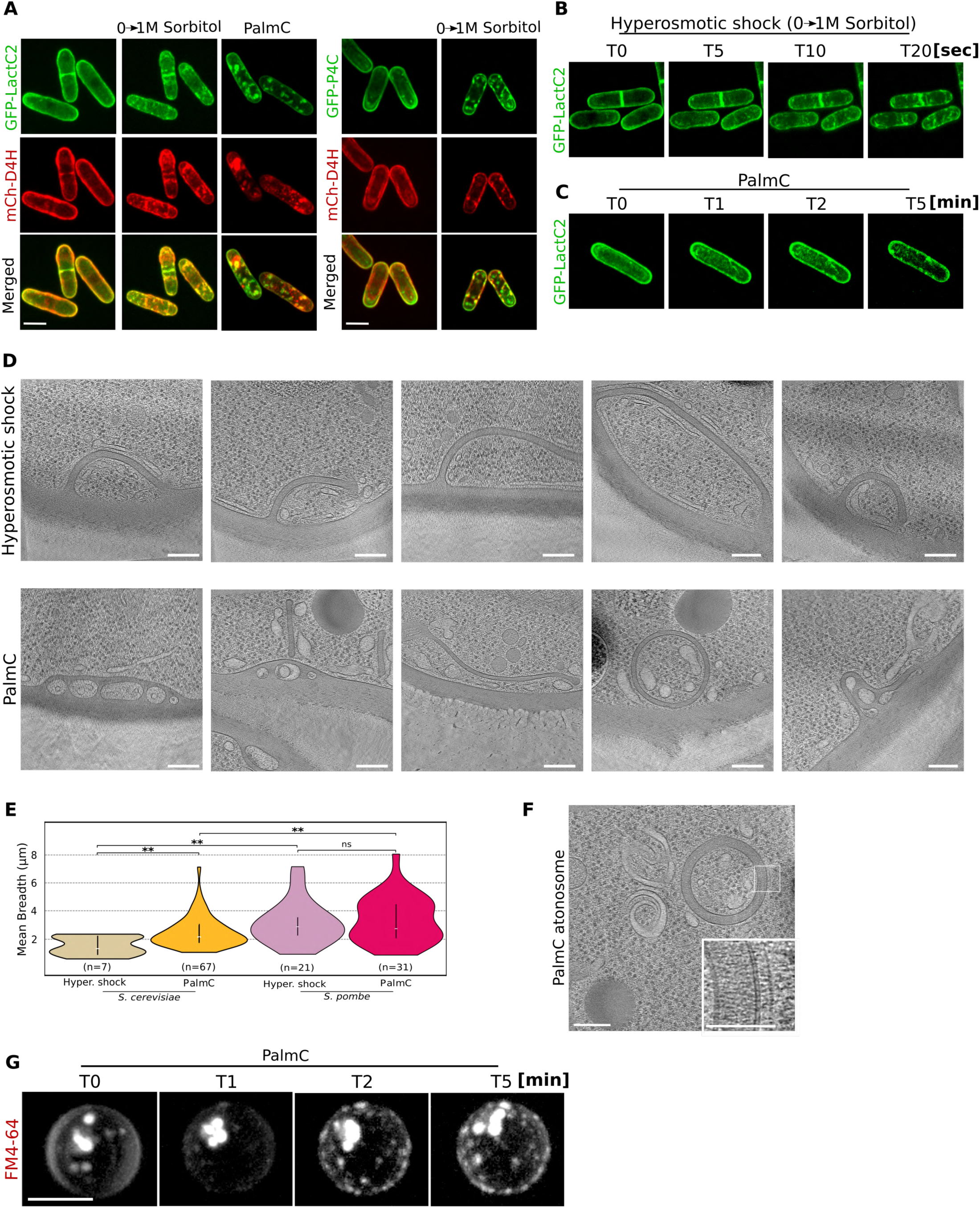
Stress-dependent dynamics and high-resolution imaging of atonosomes in unicellular cell walled-enclosed organisms. **(A)** Spinning-disk confocal microscopy of 5. *pombe* cells expressing endogenously a PS reporter (GFP-LactC2), a sterol reporter (meh-D4H) and/or a Pl4P reporter (GFP-P4C), subjected to hyperosmotic shock (1M sorbitol for 1 min) or Palme (8 µM for 5 min). Scale bar, 5 µm. **(B-C)** Live spinning-disk confocal microscopy showing rapid atonosome formation following hyperosmotic shock **(B)** or Palme treatment (8 µM) **(C)** in 5. *pombe* cells expressing endogenously a PS reporter (GFP-Lacte2). Scale bar, 5 µm. **(D)** eryo-tomographic slice of atonosomes formed under hyperosmotic shock or Palme treatment in 5. *pombe.* Scale bars, 100 nm. **(E)** Segmentation-based quantification of atonosome mean breadth in 5. *pombe,* reflecting structural complexity and comparison with 5. *cerevisiae.* "**" *p* value < 0.01; "ns" non significant; white horizontal bar, median; black vertical bar, IQR or middle 50%. **(F)** eryo-tomographic slices showing septin-like alignment along a Palme-induced atonosome surface in 5. *pombe.* Scale bars, 100 nm. Insets show magnified views of septin ring units, scale bar, 50 nm. **(G)** Live spinning-disk confocal microscopy showing rapid atonosome formation following Palme treatment (4 µM) in C. *perkinsii.* Scale bar, 5 µm. All fluorescence images are maximum-intensity projections along the z-axis

**Figure S6.**
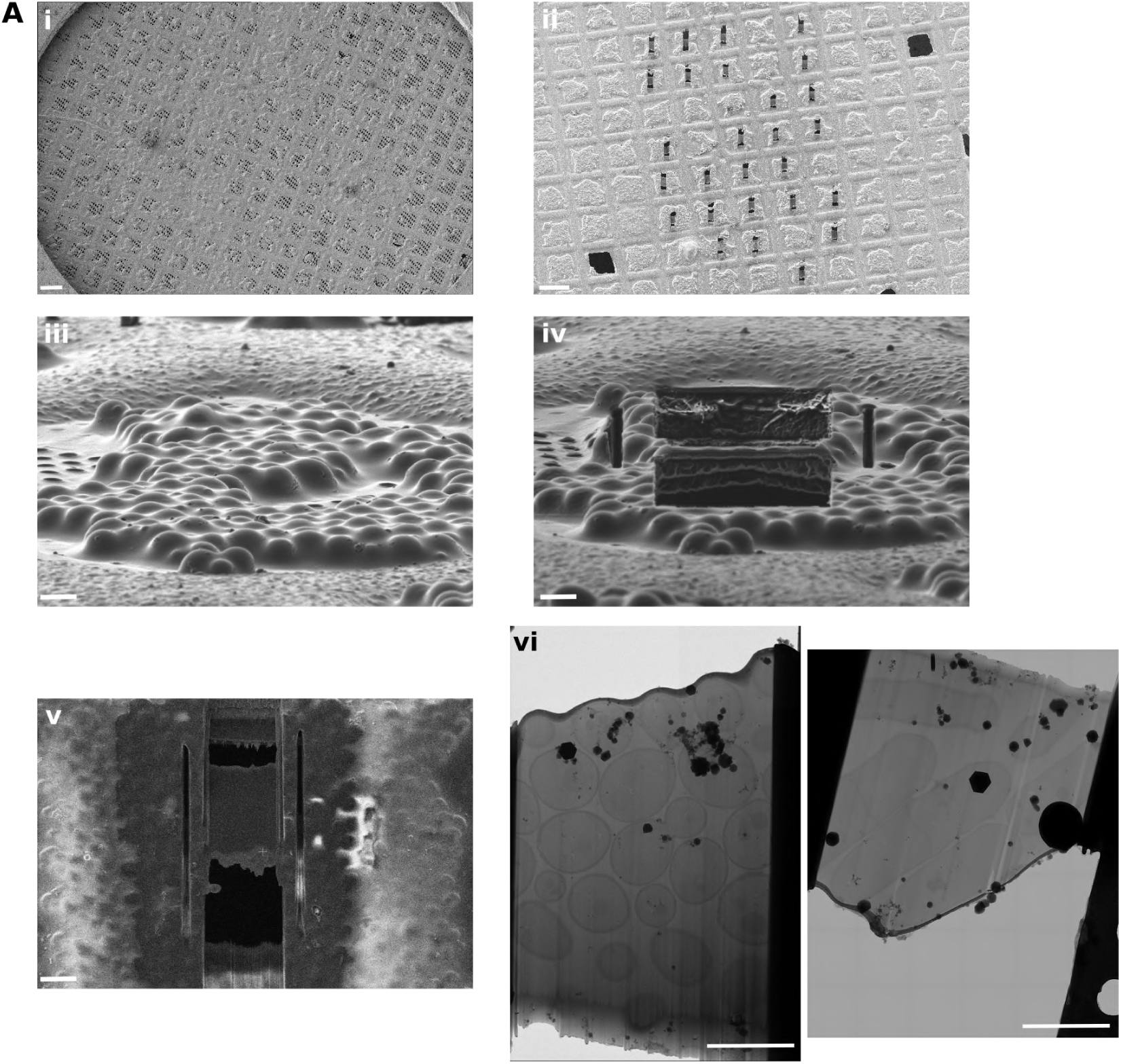
Cryo-FIB milling workflow of yeast cells. **(A)(i)** Full view of a grid covered with a homogenous layer of 5. *cerevisiae* cells. Scale bar, SO µm. **(ii)** SEM view of lamellae after the final polishing step. Scale bar, SO µm. Layer of yeast cells at the surface of the grid square before **(iii)** and after **(iv)** milling, observed from the FIB point of view. Scale bar, S µm. **(v)** Close SEM view of a representative lamella surrounded by stress relief trenches. **(vi)** TEM view of a lamella containing about 20 S. *cerevisiae* (left panel) and 7 *S.pombe* (right panel) cell sections. Scale bars, 5 µm.

**Table S1.**
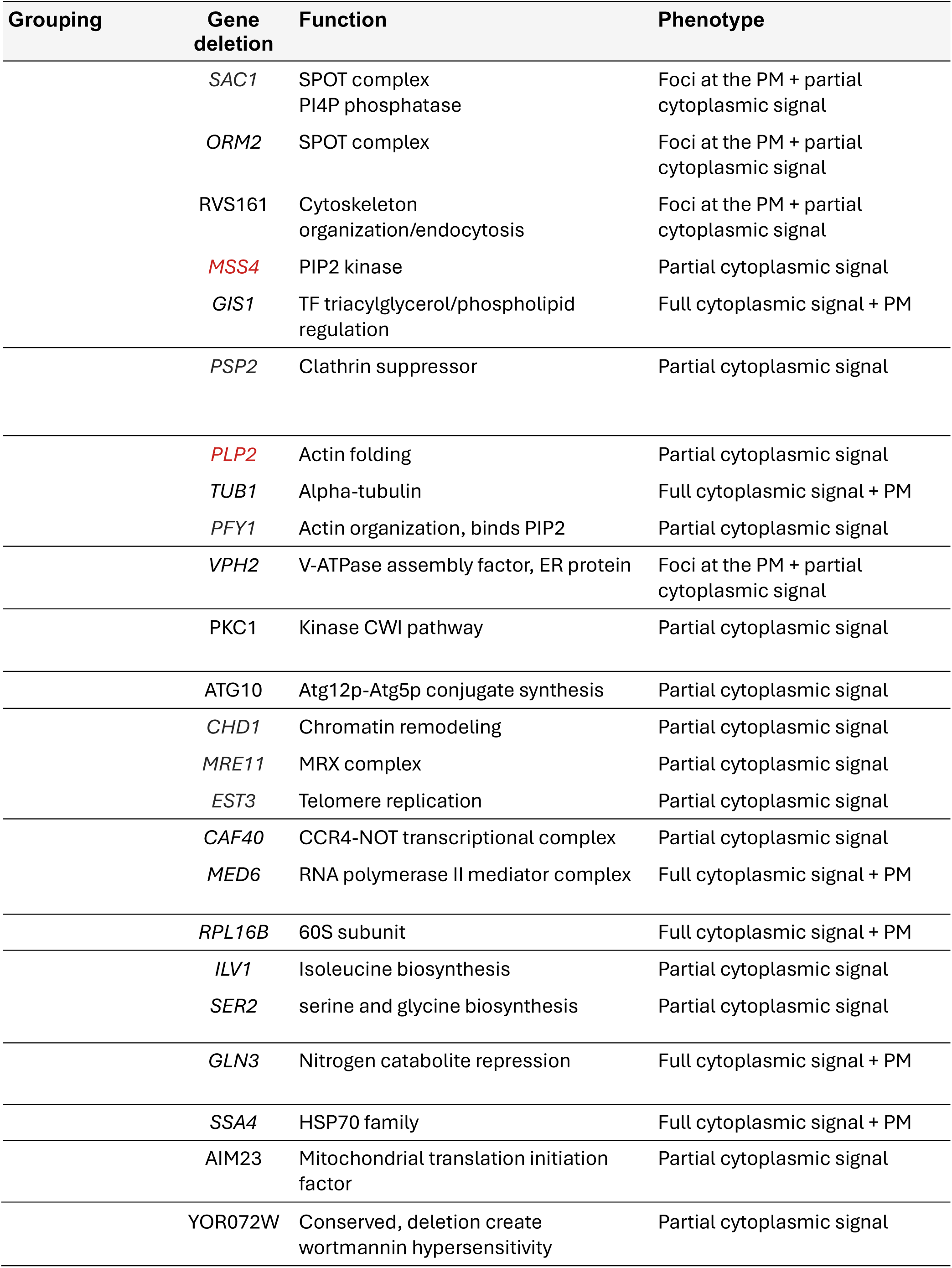
yMS2085 MATα strains carrying either a deletion of a single non-essential gene or downregulation of an essential gene (highlighted in red). All strains endogenously express fluorescent markers for PI(4,5)P₂ and phosphatidylserine (PS). Following confocal microscopy screening for plasma membrane (PM) defects, 24 strains exhibited mislocalization of one or both markers into the cytoplasm. This phenotype was either partial, with only a subset of cells displaying cytoplasmic signal for membrane markers, or total with all the cells carrying cytoplasmic localization (full). In some cells, cytoplasmic fluorescence coexisted with residual fluorescence at the PM.

## References

1. Harayama, T. & Riezman, H. Understanding the diversity of membrane lipid composition. Nat. Rev. Mol. Cell Biol. 19, 281–296 (2018).

2. Honigmann, A. & Pralle, A. Compartmentalization of the Cell Membrane. J. Mol. Biol. 428, 4739–4748 (2016).

3. Hazel, J. R. Thermal Adaptation in Biological Membranes: Is Homeoviscous Adaptation the Explanation? Annu. Rev. Physiol. 57, 19–42 (1995).

4. Kinnunen, P. K. J. Lipid Bilayers as Osmotic Response Elements. Cell. Physiol. Biochem. 10, 243–250 (2000).

5. Berchtold, D. et al. Plasma membrane stress induces relocalization of Slm proteins and activation of TORC2 to promote sphingolipid synthesis. Nat. Cell Biol. 14, 542–547 (2012).

6. Douglas, L. M. & Konopka, J. B. Fungal Membrane Organization: The Eisosome Concept. Annu. Rev. Microbiol. 68, 377–393 (2014).

7. Kefauver, J. M. et al. Cryo-EM architecture of a near-native stretch-sensitive membrane microdomain. Nature 632, 664–671 (2024).

8. Malinsky, J., Opekarová, M. & Tanner, W. The lateral compartmentation of the yeast plasma membrane. Yeast 27, 473–478 (2010).

9. Walther, T. C. et al. Eisosomes mark static sites of endocytosis. Nature 439, 998–1003 (2006).

10. Eltschinger, S. & Loewith, R. TOR Complexes and the Maintenance of Cellular Homeostasis. Trends Cell Biol. 26, 148–159 (2016).

11. Thorner, J. TOR complex 2 is a master regulator of plasma membrane homeostasis. Biochem. J. 479, 1917–1940 (2022).

12. Riggi, M. et al. Decrease in plasma membrane tension triggers PtdIns(4,5)P2 phase separation to inactivate TORC2. Nat. Cell Biol. 20, 1043–1051 (2018).

13. Phan, J. et al. Recovery of plasma membrane tension after a hyperosmotic shock. Mol. Biol. Cell 36, ar45 (2025).

14. Dupont, S., Beney, L., Ritt, J.-F., Lherminier, J. & Gervais, P. Lateral reorganization of plasma membrane is involved in the yeast resistance to severe dehydration. Biochim. Biophys. Acta BBA - Biomembr. 1798, 975–985 (2010).

15. Sakata, K.-T. et al. Coordinated regulation of TORC2 signaling by MCC/eisosome-associated proteins, Pil1 and tetraspan membrane proteins during the stress response. Mol. Microbiol. 117, 1227–1244 (2022).

16. Wang, H. X., Douglas, L. M., Aimanianda, V., Latgé, J.-P. & Konopka, J. B. The Candida albicans Sur7 protein is needed for proper synthesis of the fibrillar component of the cell wall that confers strength. Eukaryot. Cell 10, 72–80 (2011).

17. Badrane, H. et al. Rapid Redistribution of Phosphatidylinositol-(4,5)-Bisphosphate and Septins during the Candida albicans Response to Caspofungin. Antimicrob. Agents Chemother. 56, 4614–4624 (2012).

18. Stefan, C. J., Audhya, A. & Emr, S. D. The Yeast Synaptojanin-like Proteins Control the Cellular Distribution of Phosphatidylinositol (4,5)-Bisphosphate. Mol. Biol. Cell 13, 542–557 (2002).

19. Stolz, L. E., Huynh, C. V., Thorner, J. & York, J. D. Identification and characterization of an essential family of inositol polyphosphate 5-phosphatases (INP51, INP52 and INP53 gene products) in the yeast Saccharomyces cerevisiae. Genetics 148, 1715–1729 (1998).

20. Bharat, T. A. M., Hoffmann, P. C. & Kukulski, W. Correlative Microscopy of Vitreous Sections Provides Insights into BAR-Domain Organization In Situ. Struct. Englan*d1993* 26, 879–886.e3 (2018).

21. Andersen, M. H., Graversen, H., Fedosov, S. N., Petersen, T. E. & Rasmussen, J. T. Functional analyses of two cellular binding domains of bovine lactadherin. Biochemistry 39, 6200–6206 (2000).

22. Kavran, J. M. et al. Specificity and promiscuity in phosphoinositide binding by pleckstrin homology domains. J. Biol. Chem. 273, 30497–30508 (1998).

23. Tettamanti, M. G. et al. A dynamic feedback loop between retrograde sterol transport and TORC2 controls adaptation of the plasma membrane to stress. EMBO J. 44, 7541–7564 (2025).

24. Moreira, K. E., Walther, T. C., Aguilar, P. S. & Walter, P. Pil1 Controls Eisosome Biogenesis. Mol. Biol. Cell 20, 809–818 (2009).

25. Niles, B. J., Mogri, H., Hill, A., Vlahakis, A. & Powers, T. Plasma membrane recruitment and activation of the AGC kinase Ypk1 is mediated by target of rapamycin complex 2 (TORC2) and its effector proteins Slm1 and Slm2. Proc. Natl. Acad. Sci. U. S. A. 109, 1536–1541 (2012).

26. Breslow, D. K. et al. A comprehensive strategy enabling high-resolution functional analysis of the yeast genome. Nat. Methods 5, 711–718 (2008).

27. Giaever, G. et al. Functional profiling of the Saccharomyces cerevisiae genome. Nature 418, 387–391 (2002).

28. Schäfer, J.-H. et al. Structure of the ceramide-bound SPOTS complex. Nat. Commun. 14, 6196 (2023).

29. Shimobayashi, M., Oppliger, W., Moes, S., Jenö, P. & Hall, M. N. TORC1-regulated protein kinase Npr1 phosphorylates Orm to stimulate complex sphingolipid synthesis. Mol. Biol. Cell 24, 870–881 (2013).

30. Liu, K., Kong, L., Graham, D. B., Carey, K. L. & Xavier, R. J. SAC1 regulates autophagosomal phosphatidylinositol-4-phosphate for xenophagy-directed bacterial clearance. Cell Rep. 36, 109434 (2021).

31. Zewe, J. P., Wills, R. C., Sangappa, S., Goulden, B. D. & Hammond, G. R. SAC1 degrades its lipid substrate PtdIns4P in the endoplasmic reticulum to maintain a steep chemical gradient with donor membranes. eLife 7, e35588 (2018).

32. Bhattacharya, S., Esquivel, B. D. & White, T. C. Overexpression or Deletion of Ergosterol Biosynthesis Genes Alters Doubling Time, Response to Stress Agents, and Drug Susceptibility in Saccharomyces cerevisiae. mBio 9, e01291–18 (2018).

33. Abe, F. & Hiraki, T. Mechanistic role of ergosterol in membrane rigidity and cycloheximide resistance in *Saccharomyces cerevisiae*. Biochim. Biophys. Acta BBA - Biomembr. 1788, 743–752 (2009).

34. Valachovic, M. et al. Cumulative Mutations Affecting Sterol Biosynthesis in the Yeast Saccharomyces cerevisiae Result in Synthetic Lethality That Is Suppressed by Alterations in Sphingolipid Profiles. Genetics 173, 1893–1908 (2006).

35. Aresta-Branco, F. et al. Gel Domains in the Plasma Membrane of Saccharomyces cerevisiae: HIGHLY ORDERED, ERGOSTEROL-FREE, AND SPHINGOLIPID-ENRICHED LIPID RAFTS *. J. Biol. Chem. 286, 5043–5054 (2011).

36. Kapteyn, J. C., Van Den Ende, H. & Klis, F. M. The contribution of cell wall proteins to the organization of the yeast cell wall. Biochim. Biophys. Acta BBA - Gen. Subj. 1426, 373–383 (1999).

37. Klis, F. M., Mol, P., Hellingwerf, K. & Brul, S. Dynamics of cell wall structure in Saccharomyces cerevisiae. FEMS Microbiol. Rev. 26, 239–256 (2002).

38. Costa-de-Oliveira, S. et al. Determination of chitin content in fungal cell wall: An alternative flow cytometric method. Cytometry A 83A, 324–328 (2013).

39. de Assis, L. J. et al. Nature of β-1,3-Glucan-Exposing Features on Candida albicans Cell Wall and Their Modulation. mBio 13, e02605–22 (2022).

40. Wood, V. Schizosaccharomyces pombe comparative genomics; from sequence to systems. in Comparative Genomics: Using Fungi as Models (eds Sunnerhagen, P. & Piskur, J.) 233–285 (Springer, Berlin, Heidelberg, 2006). doi:10.1007/4735_97.

41. Maekawa, M., Yang, Y. & Fairn, G. D. Perfringolysin O Theta Toxin as a Tool to Monitor the Distribution and Inhomogeneity of Cholesterol in Cellular Membranes. Toxins 8, 67 (2016).

42. Luo, X. et al. Structure of the Legionella Virulence Factor, SidC Reveals a Unique PI(4)P-Specific Binding Domain Essential for Its Targeting to the Bacterial Phagosome. PLOS Pathog. 11, e1004965 (2015).

43. Grau-Bové, X. et al. Dynamics of genomic innovation in the unicellular ancestry of animals. eLife 6, e26036 (2017).

44. Shah, H. et al. Life-cycle-coupled evolution of mitosis in close relatives of animals. Nature 630, 116–122 (2024).

45. Olivetta, M., Bhickta, C., Chiaruttini, N., Burns, J. & Dudin, O. A multicellular developmental program in a close animal relative. Nature 635, 382–389 (2024).

46. Sambre, P. D., Ho, J. C. S. & Parikh, A. N. Intravesicular Solute Delivery and Surface Area Regulation in Giant Unilamellar Vesicles Driven by Cycles of Osmotic Stresses. J. Am. Chem. Soc. 146, 3250–3261 (2024).

47. Arumugam, S. & Bassereau, P. Membrane nanodomains: contribution of curvature and interaction with proteins and cytoskeleton. Essays Biochem. 57, 109–119 (2015).

48. Fessler, M. B. & Parks, J. S. Intracellular Lipid Flux and Membrane Microdomains as Organizing Principles in Inflammatory Cell Signaling. J. Immunol. 187, 1529–1535 (2011).

49. Lou, H.-Y., Zhao, W., Zeng, Y. & Cui, B. The Role of Membrane Curvature in Nanoscale Topography-Induced Intracellular Signaling. Acc. Chem. Res. 51, 1046–1053 (2018).

50. Mollinedo, F. Lipid raft involvement in yeast cell growth and death. Front. Oncol. 2, 140 (2012).

51. Bertin, A. et al. Phosphatidylinositol-4,5-bisphosphate promotes budding yeast septin filament assembly and organization. J. Mol. Biol. 404, 711–731 (2010).

52. Szuba, A. et al. Membrane binding controls ordered self-assembly of animal septins. eLife 10, e63349 (2021).

53. de Nadal, E. & Posas, F. The HOG pathway and the regulation of osmoadaptive responses in yeast. FEMS Yeast Res. 22, foac013 (2022).

54. Sakata, K. et al. Slm1, an F-BAR and PH domain-containing protein, regulates heat-induced turnover of nutrient transporter to control plasma-membrane quality. FEBS J. n/a, (2026).

55. Rodríguez-Escudero, I., Fernández-Acero, T., Cid, V. J. & Molina, M. Heterologous mammalian Akt disrupts plasma membrane homeostasis by taking over TORC2 signaling in Saccharomyces cerevisiae. Sci. Rep. 8, 7732 (2018).

56. Feyder, S., De Craene, J.-O., Bär, S., Bertazzi, D. L. & Friant, S. Membrane Trafficking in the Yeast Saccharomyces cerevisiae Model. Int. J. Mol. Sci. 16, 1509–1525 (2015).

57. Armstrong, J. Yeast vacuoles: more than a model lysosome. Trends Cell Biol. 20, 580–585 (2010).

58. Li, S. C. & Kane, P. M. The Yeast Lysosome-like Vacuole: Endpoint and Crossroads. Biochim. Biophys. Acta 1793, 650–663 (2009).

59. Costa, V. & Teixeira, V. Vacuolar ATPase-mediated regulation of neutral lipid dynamics: Insights into lipid droplet homeostasis and stress response mechanisms. Biochim. Biophys. Acta Mol. Cell Biol. Lipids 1869, 159465 (2024).

60. Mazheika, I. S., Psurtseva, N. V. & Kamzolkina, O. V. Lomasomes and Other Fungal Plasma Membrane Macroinvaginations Have a Tubular and Lamellar Genesis. J. Fungi 8, 1316 (2022).

61. Daraspe, J., Bellani, E., De Bellis, D., Genoud, C. & Geldner, N. Targeted imaging of specialized plant cell walls by improved cryo-CLEM and cryo-electron tomography. Nat. Commun. 16, 11354 (2025).

62. Dakhel, A. et al. Zombosomes are anucleated cell couriers that spread α-synuclein pathology. Cell Rep. 45, (2026).

63. Jeppesen, D. K. et al. Blebbisomes are large, organelle-rich extracellular vesicles with cell-like properties. Nat. Cell Biol. 27, 438–448 (2025).

64. Marek, M., Vincenzetti, V. & Martin, S. G. Sterol biosensor reveals LAM-family Ltc1-dependent sterol flow to endosomes upon Arp2/3 inhibition. J. Cell Biol. 219, e202001147 (2020).

65. Asano, S., Engel, B. D. & Baumeister, W. In Situ Cryo-Electron Tomography: A Post-Reductionist Approach to Structural Biology. J. Mol. Biol. 428, 332–343 (2016).

66. Rigort, A. et al. Focused ion beam micromachining of eukaryotic cells for cryoelectron tomography. Proc. Natl. Acad. Sci. 109, 4449–4454 (2012).

67. Turoňová, B. et al. Benchmarking tomographic acquisition schemes for high-resolution structural biology. Nat. Commun. 2020 111 11, 1–9 (2020).

68. Zheng, S. Q. et al. MotionCor2: Anisotropic correction of beam-induced motion for improved cryo-electron microscopy. Nat. Methods 14, 331–332 (2017).

69. Mastronarde, D. Tomographic Reconstruction with the IMOD Software Package. Microsc. Microanal. 12, 178–179 (2006).

70. Galaz-Montoya, J. G., Flanagan, J., Schmid, M. F. & Ludtke, S. J. Single particle tomography in EMAN2. J. Struct. Biol. 190, 279–290 (2015).

71. Lamm, L. et al. MemBrain: A Deep Learning-aided Pipeline for Automated Detection of Membrane Proteins in Cryo-electron Tomograms. 2022.03.01.480844 Preprint at 10.1101/2022.03.01.480844 (2022).

72. Lamm, L. et al. MemBrain v2: an end-to-end tool for the analysis of membranes in cryo-electron tomography. 2024.01.05.574336 Preprint at 10.1101/2024.01.05.574336 (2025).

73. Goddard, T. D. et al. UCSF ChimeraX: Meeting modern challenges in visualization and analysis. Protein Sci. Publ. Protein Soc. 27, 14–25 (2018).

74. Sofroniew, N. et al. napari: a multi-dimensional image viewer for Python. 10.5281/zenodo.20709435 (2026) doi:10.5281/zenodo.20709435.

75. Schindelin, J., et al. Fiji: an open-source platform for biological-image analysis. Nat. Methods 9, 676–682 (2012).

76. Miura, K. Bleach correction ImageJ plugin for compensating the photobleaching of time-lapse sequences. Preprint at 10.12688/f1000research.27171.1 (2020).

77. Laine, R. F. et al. NanoJ: a high-performance open-source super-resolution microscopy toolbox. J. Phys. Appl. Phys. 52, 163001 (2019).

78. Pylvänäinen, J. W. et al. Fast4DReg – fast registration of 4D microscopy datasets. J. Cell Sci. 136, jcs260728 (2023).

79. Legland, D., Arganda-Carreras, I. & Andrey, P. MorphoLibJ: integrated library and plugins for mathematical morphology with ImageJ. Bioinformatics 32, 3532–3534 (2016).

80. Weill, U. et al. Genome-wide SWAp-Tag yeast libraries for proteome exploration. Nat. Methods 15, 617–622 (2018).

81. Yan Tong, A. H. & Boone, C. Synthetic Genetic Array Analysis in Saccharomyces cerevisiae. in Yeast Protocol (ed. Xiao, W.) 171–191 (Humana Press, Totowa, NJ, 2006). doi:10.1385/1-59259-958-3:171.

82. Schneider, C. A., Rasband, W. S. & Eliceiri, K. W. NIH Image to ImageJ: 25 years of image analysis. Nat. Methods 9, 671–675 (2012).

83. Zahn-Zabal, M., Dessimoz, C. & Glover, N. M. Identifying orthologs with OMA: A primer. F1000Research 9, 27 (2020).

84. Waterhouse, A. M., Procter, J. B., Martin, D. M. A., Clamp, M. & Barton, G. J. Jalview Version 2—a multiple sequence alignment editor and analysis workbench. Bioinformatics 25, 1189–1191 (2009).

