## Supplementary material for "Atonosomes, compartments involved in membrane tension decrease": Movie legend

**Commented [RL1]:** Need to indicate author affiliations

### Movie legends

**Commented [EB2]:** Elda Bauda
Comment Margot : Would maybe remove and just insist on what you think they are instead of what they're not. I find "simple invaginations" a bit of an oversimplification of the previous findings in the literature and in particular, the model from the 2025 Babst paper already explicitly propose a different topology than invaginations. "Here we propose tonosomes as a unifying identity for a class of PM-derived compartments arising across contexts of acute and chronic membrane homeostatic stress (or: acute and chronic tensile and lipid homeostatic stress), leveraging unprecedented high res ultrastructural imaging as a scaffold to integrate previously disparate observations reported under various names."

S1 . Cryo tomogram and segmentation of tonosome shown in Fig. 1A.

S2a. Live confocal spinning disk imaging of tonosome formation upon hyperosmotic shock (1M sorbitol), monitored using a PI(4,5)$P_2$ probe (mCh-2xPH). Time is indicated in minutes:seconds.

S2b. Live confocal spinning disk imaging of tonosome formation upon PalmC treatment, monitored using a PI(4,5)$P_2$ probe (mCh-2xPH). Time is indicated in minutes:seconds.

S3a. Live confocal spinning disk imaging of septin recruitment (Cdc10-GFP) to tonosomes (PI(4,5)$P_2$ probe (mCh-2xPH ) upon hyperosmotic shock (1M sorbitol). Time is indicated in minutes:seconds.

S3b. Live confocal spinning disk imaging of septin recruitment (Cdc10-GFP) to tonosomes (PI(4,5)$P_2$ probe (mCh-2xPH ) upon PalmC treatment. Time is indicated in minutes:seconds.

S4. Cryo tomogram of constitutive tonosome shown in Fig. 3A.

S5. Serial sectioning of constitutive tonosome in *pil1Δlsp1Δ* cells. Each frame of the movie represent a successive 60 nm thick section. CW appears white.

S6a. Live confocal spinning disk imaging showing the reaction of constitutive thekosomes in *pil1Δlsp1Δ6tsp Δ cells* upon hyperosmotic shock (1M sorbitol), monitored using a PI(4,5)$P_2$ probe (mCh-2xPH). Time is indicated in minutes:seconds.

S6b. Live confocal spinning disk imaging showing the reaction of constitutive thekosomes in *pil1Δlsp1Δ6tsp Δ cells* upon PalmC treatment, monitored using a PI(4,5)$P_2$ probe (mCh-2xPH). Time is indicated in minutes:seconds.

S7a. Live confocal spinning disk imaging of tonosome formation in p*il1Δlsp1Δslm1/2 Δ6tspΔ* cells upon hyperosmotic shock (1M sorbitol), monitored using a PI(4,5)$P_2$ probe (mCh-2xPH). Time is indicated in minutes:seconds.

S7b. Live confocal spinning disk imaging of tonosome formation in p*il1Δlsp1Δslm1/2 Δ6tspΔ* cells upon PalmC treatment, monitored using a PI(4,5)$P_2$ probe (mCh-2xPH). Time is indicated in minutes:seconds.

S8a. Live confocal spinning disk imaging the resorption of hyperosmotic shock-induced tonosomes during hypoosmotic shock (1 M → 0 M sorbitol) in WT cells, imaged using the PI(4,5)P2 probe (mCherry-2xPH). Time is indicated in minutes:seconds.

S8b. Live confocal spinning disk imaging the resorption of PalmC-induced tonosomes during hypoosmotic shock (1 M → 0 M sorbitol) in WT cells, imaged using the PI(4,5)P2 probe (mCherry-2xPH). Time is indicated in minutes:seconds.

S8c. Live confocal spinning disk imaging of constitutive tomosomes in in p*il1Δlsp1Δ* cells during hypoosmotic shock (1 M → 0 M sorbitol), visualized using the PI(4,5)P2 probe (mCherry-2xPH). Time is indicated in minutes:seconds.

S8c. Live confocal spinning disk imaging of constitutive tomosomes in p*il1Δlsp1Δ6tspΔ* cells during hypoosmotic shock (1 M → 0 M sorbitol), visualized using the PI(4,5)P2 probe (mCherry-2xPH). Time is indicated in minutes:seconds

S9. Live confocal spinning disk imaging of protoplasts submitted to PalmC treatment, monitored using the PI(4,5)P2 probe (mCherry-2xPH) and pil1-GFP. Time is indicated in minutes:seconds

S10a. Live confocal spinning disk imaging of tonosome formation in *S. pombe* upon hyperosmotic shock (1M sorbitol), monitored using a PI4P probe (xxxx) and a sterol probe (mCh-D4H). Time is indicated in minutes:seconds.

S10b. Live confocal spinning disk imaging of tonosome formation in *S. pombe* upon PalmC treatment , monitored using a PI4P probe (xxxx) and a sterol probe (mCh-D4H). Time is indicated in minutes:seconds

S11. Cryo tomogram of tonosome in *S. pombe* shown in Fig. 6A.

S12. Live confocal spinning disk imaging of tonosome formation in *C. perkinsii* upon PalmC treatment , monitored using FM4-64. Time is indicated in minutes:seconds
